# Integrated single-cell transcriptomics and epigenomics reveals strong germinal center-associated etiology of autoimmune risk loci

**DOI:** 10.1101/2021.03.16.435578

**Authors:** Hamish W King, Kristen L Wells, Zohar Shipony, Arwa S Kathiria, Lisa E Wagar, Caleb Lareau, Nara Orban, Robson Capasso, Mark M Davis, Lars M Steinmetz, Louisa K James, William J Greenleaf

## Abstract

The germinal center (GC) response is critical for both effective adaptive immunity and establishing peripheral tolerance by limiting auto-reactive B cells. Dysfunction in these processes can lead to defects in immune response to pathogens or contribute to autoimmune disease. To understand the gene regulatory principles underlying the GC response, we generated a single-cell transcriptomic and epigenomic atlas of the human tonsil, a widely studied and representative lymphoid tissue. We characterize diverse immune cell subsets and build a trajectory of dynamic gene expression and transcription factor activity during B cell activation, GC formation, and plasma cell differentiation. We subsequently leverage cell type-specific transcriptomic and epigenomic maps to interpret potential regulatory impact of genetic variants implicated in autoimmunity, revealing that many exhibit their greatest regulatory potential in GC cell populations. Together, these analyses provide a powerful new cell type-resolved resource for the interpretation of cellular and genetic causes underpinning autoimmune disease.

**One sentence summary:** Single-cell chromatin accessibility landscapes of immune cell subsets reveal regulatory potential of autoimmune-associated genetic variants during the germinal center response.

## Introduction

Autoimmune diseases result from a loss of tolerance to otherwise harmless endogenous or exogenous antigens, in part as a consequence of dysregulation in the selection, differentiation, or function of immune cells. The propensity for such immune cell dysfunction can be potentiated by specific inherited genetic variants, as identified through genome wide association studies (GWAS). However, the majority of GWAS genetic variants reside in non-coding regions of the genome, and the identification of risk-associated genetic variants alone does not identify the cellular populations likely affected by the variant. Recent progress has been made linking autoimmune-associated genetic variants to immune cell type-specific gene regulation by examining functional epigenomic measures like chromatin accessibility, histone acetylation and/or chromatin topology, especially in activated immune cell states of immune subsets (1–3). However, such analysis remains incomplete due to limited mapping of important yet transient subpopulations of cells that exist in diverse immune organ contexts.

The development and commitment of different immune cell lineages occurs in primary lymphoid organs such as the bone marrow and thymus. Following lineage commitment and egress from these organs, adaptive immune cells can undergo additional maturation and differentiation in secondary lymphoid organs such as the spleen, lymph nodes and tonsils to generate T cell-mediated immunity and B cell-dependent antibody responses (4). The latter in particular is predominantly dependent on the formation of the germinal center (GC) response. This requires MHCII-dependent presentation of antigen-derived peptides by dendritic cells that can be recognized by naïve CD4^+^ T cells, leading to their differentiation into T follicular helper (Tfh) cells. Tfh are vital to support activated B cells to form GC reactions, undergo somatic hypermutation and affinity maturation of their antibody genes before differentiating into plasma cells or memory B cells.

Mechanisms that ensure immune tolerance to self-antigen target autoreactive B cell clones during both early development in the bone marrow (central tolerance) and de novo generation in GC responses in secondary lymphoid organs (peripheral tolerance). Autoantibodies are a feature of many systemic autoimmune diseases, and numerous studies have found that autoantibodies can bear somatic hypermutation and class switching signatures indicative of GC-derived B cell populations (5), pointing to defects in peripheral tolerance. Because these tissues and GC-associated immune cell populations are directly involved in establishing both peripheral tolerance and forming effective adaptive immune responses, mapping the regulatory potential of autoimmune-associated genetic variants in these dynamic populations will enable the interpretation of how these variants may contribute to autoimmunity.

Here we apply single-cell transcriptomics (scRNA-seq), surface-protein profiling (scADT-seq), and epigenomics (scATAC-seq) to map the cellular states and gene regulatory networks of immune cells from the human tonsil, a model secondary lymphoid organ (Fig1-2). By integrating gene expression and chromatin accessibility across 37 immune cell populations spanning bone marrow, peripheral blood, and tonsils, we predict putative target genes of fine-mapped autoimmune-associated genetic variants and discover extensive GC-specific regulatory potential (Fig3-4), including at loci of major GC regulators such as IL21, IL21R/IL4R and BCL6 (Fig5), as well as two genes required for MBC fate commitment, POU2AF1 and HHEX (Fig6). Our integrative analyses ultimately provide new insights into the cellular and genetic etiology of autoimmune-associated genetic variants and generate a framework to functionally dissect their potential in the maintenance of peripheral tolerance and the generation of adaptive immunity.

**Figure 1.**
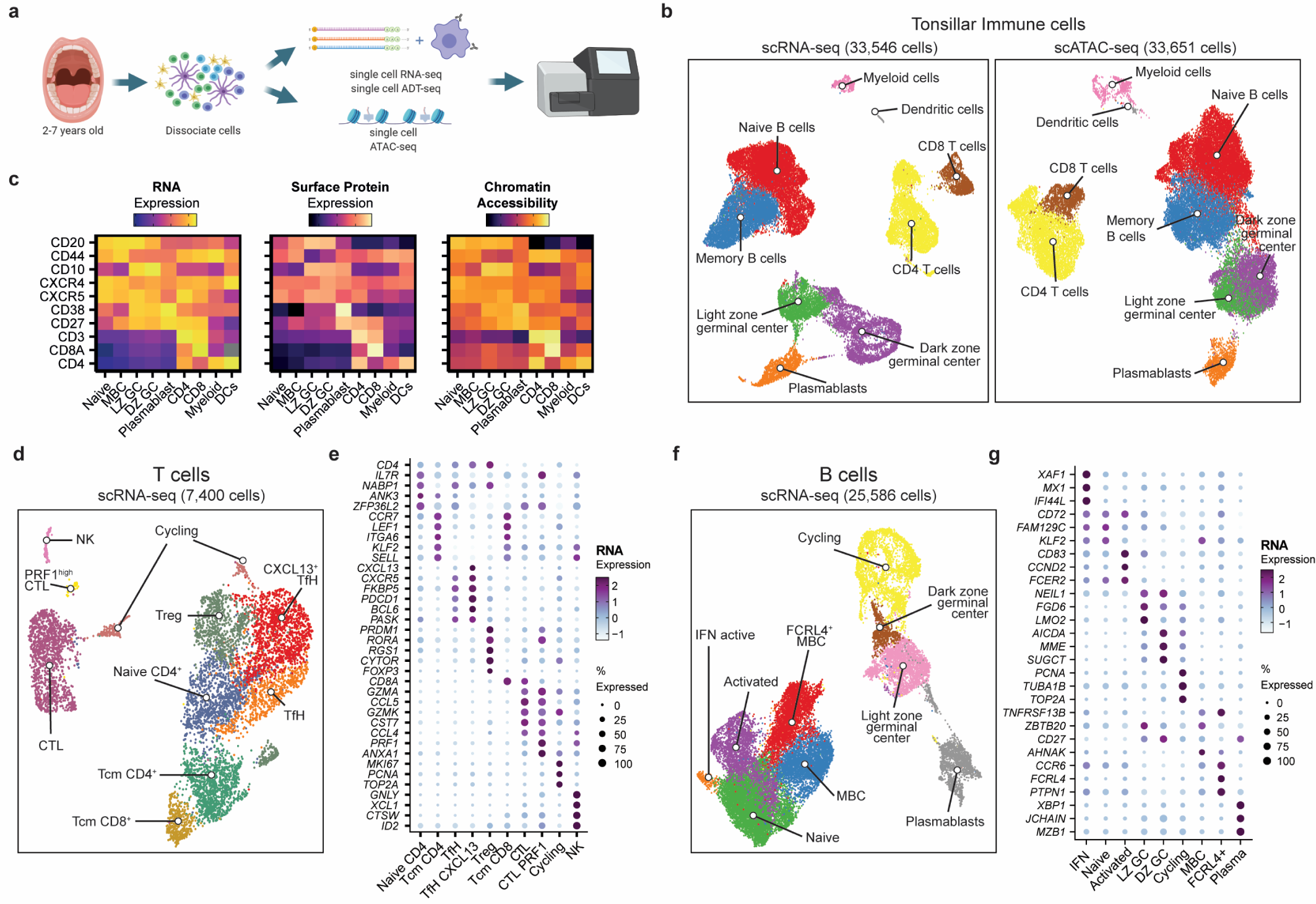
Single-cell mapping of immune cell subsets in human tonsils. a) Experimental strategy for single-cell transcriptomics, surface marker expression, and chromatin accessibility of immune cells from pediatric tonsils. b) UMAP of tonsillar immune scRNA-seq data (left; 3 donors) and scATAC-seq data (right; 7 donors). c) Heatmap comparing gene expression, surface protein, and chromatin accessibility across immune cell types. d) UMAP of T cell sub-populations in the tonsillar immune scRNA-seq data in B). NK = natural killer, CTL = cytotoxic lymphocyte, Treg = regulatory T cell, TfH = T follicular helper cell, Tcm = T central memory. e) Mean expression of key marker genes for T cell sub-populations by scRNA-seq. Frequency of cells for which each gene is detected is denoted by size of the dots. f) UMAP of B cell sub-populations in the tonsillar immune scRNA-seq data in B). MBC = memory B cell, LZ GC = light zone germinal center, DZ GC = dark zone germinal center; IFN = interferon. g) Mean expression of key marker genes for B cell sub-populations by scRNA-seq. Frequency of cells for which each gene is detected is denoted by size of the dots.

## Results

### Single-cell transcriptomics and epigenomics of a model human secondary lymphoid organ to define immune cell states

To map the diverse immune cell states of the adaptive immune response in human secondary lymphoid organs, and the gene regulatory elements active in these different populations, we performed high throughput single-cell RNA sequencing (scRNA-seq) coupled with antibody-derived tags (scADT-seq) for twelve surface protein markers on unsorted tonsillar immune cells (Fig1a-c, Table S1). In parallel, we performed single-cell assay for transposase-accessible chromatin using sequencing (6) (scATAC-seq) to profile active chromatin regulatory elements (Fig1a-c; Fig 2 for more detailed analysis). We first annotated 9 broad populations based on their surface protein and RNA levels of known markers (Fig1b) and observed good concurrence between RNA, surface protein expression, and chromatin accessibility of key marker genes and the frequency of different cell types (Fig1c, S1-2, Table S2). We observed decreasing B cell frequencies with age by scRNA-seq (FigS3a), and CyTOF profiling of pediatric and adult tonsils revealed clear reductions in GC-specific B and T cell populations with age (FigS3b-d), consistent with reduced GC activity in older individuals (7, 8). As the GC is the site of major cell fate decisions during adaptive immune responses, this demonstrates the need to profile pediatric and/or immunologically relevant (e.g. after vaccination or infection) lymphoid tissue, in contrast to peripheral blood-derived immune populations or lymphoid tissue from older individuals that lack these populations.

**Figure 2.**
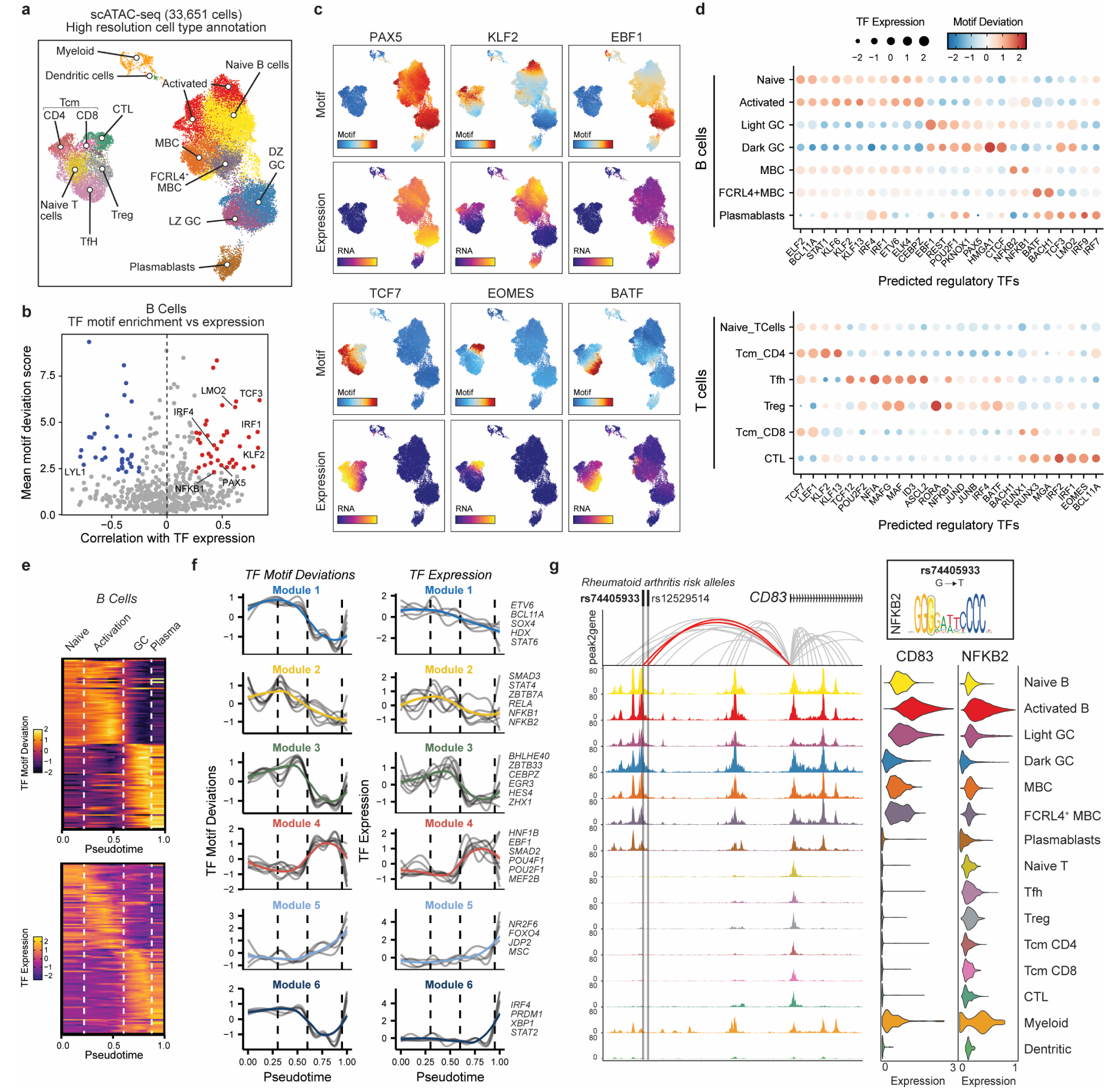
Tonsillar immune cell type-specific transcription factor regulatory activity. a) UMAP of tonsillar immune scATAC-seq with high resolution annotation of immune cell types. b) Correlation of TF motif deviation (enrichment) scores with TF expression (x axis) compared to TF motif deviation scores (y axis) to predict positive TF regulators across B cell populations. c) Motif deviation scores (top panels) and RNA expression (bottom) for exemplar TFs. d) Motif deviation scores for transcription factors (expressed in >25% cells in at least one cell type cluster). Mean gene expression is depicted by dot size. e) Pseudotemporal reconstruction of B cell activation, GC entry and plasmablast differentiation using scATAC-seq. Dotted lines highlight major transition points between cell types. Top; TF motif deviations. Bottom; TF gene expression. f) Grouped patterns of TF motif deviations (left) and TF gene expression (right) through B cell pseudotemporal reconstruction shown in (e). Colored line represents mean of all TFs per group (listed on right). g) Genomic snapshot of tonsillar immune cell scATAC-seq tracks at CD83 locus, highlighting rheumatoid arthritis-associated SNPs rs74405933 and rs12529514 and peak2gene linkages. rs74405933 falls within an NFKB2 predicted binding site (G→T). scRNA-seq expression of CD83 and NFKB2 are shown to the right.

We next annotated B or T lymphocyte sub-populations at a higher resolution using our scRNA-seq dataset (Fig1d-g, TableS3-4). Within the T cell lineage, we identified naïve and central memory T (Tcm) cells, cytotoxic lymphocytes (CTL), NK cells, regulatory T cells (Treg) and two populations of Tfh cells, with one population expressing high levels of CXCL13 and likely representing GC Tfh (Fig1d-e). We also defined clusters with gene expression markers for many expected B cell populations, including naïve, activated, memory, tissue-resident FCRL4+ memory, GC (light zone and dark zone) B cells, as well as plasmablasts (Fig1f-g). We also found a small and unexpected cluster of B cells expressing markers of type I interferon response genes such as IFI44L, XAF1, and MX1 (Fig1f-g) that are known to be up-regulated after early stages of vaccination (9). These annotations broadly agreed with recent single-cell studies of lymphocytes in pediatric tonsils and adult lymph node (10, 11).

### Mapping chromatin accessibility and transcription factor activity in tonsillar immune subsets

Our high-resolution annotation of immune cell populations by scRNA-seq (Fig 1) allowed us to more comprehensively annotate our scATAC datasets (Fig2a; see Methods for details) (12). We limited our annotations of the chromatin accessibility maps to 14 cell populations to maximize coverage and representation of cell type-specific peaks in subsequent analyses. We identified naive, activated, memory, FCRL4+ memory and GC (light zone and dark zone) B cell subsets, as well as plasmablasts, Tfh, Treg, naive, central memory and cytotoxic T cells, and two smaller clusters representing a combination of monocytes, macrophages and dendritic cells (Fig2a). We found a strong correspondence between cluster identities and cell type-specific markers used in both scATAC-seq and scRNA-seq annotation of our datasets (FigS1-2). Cells at different stages of the cell cycle, such as proliferating dark zone GC B cells, were difficult to distinguish based on their chromatin accessibility profiles, as we and others have observed few qualitative differences in chromatin accessibility profiles between mitotic and interphase cells (13, 14). Across these 14 tonsillar immune cell populations, we provide a comprehensive resource of cell type-specific gene regulatory elements in this model secondary lymphoid organ (FigS4, TableS5-6), including at the immunoglobulin heavy chain locus (FigS4c).

Lymphocyte activation, maturation, and differentiation are underpinned by transcriptional networks under the control of sequence-specific transcription factors (TFs). To understand the regulatory potential of different TFs in vivo we correlated the expression of TFs with the chromatin accessibility of their target motif sequences in both defined lymphocyte populations (Fig2b-d) and a pseudotemporal reconstruction of B cell activation, the GC reaction and plasmablast differentiation (Fig2e-f). This analysis revealed shared regulatory TF activities between similar cell states, such as those active in naive, activated and memory B cells (KLF2, BCL11A, ELF2, ETV6, ELK4) or GC B cells (EBF1, REST, POU2F1, PKNOX1) (Fig2c-d). We also found highly cell type-specific activities such as for EOMES, IRF1/2 and RUNX1/3 in cytotoxic lymphocytes, and ID3, ASCL2, NFIA and TCF12 in Tfh cells (Fig2c-d).

Meanwhile, our pseudotemporal ordering of B cell activation revealed distinct modules of TF activity and expression before commitment to the GC state (Fig2e-f), including an early activity of NFκB family members (Module 2; REL, RELA, NFKB1, NFKB2). We identified a NFκB/RELA binding site predicted to be disrupted by a rheumatoid arthritis (RA)-associated SNP (rs74405933; G→T) that is strongly linked with CD83 expression (Fig2g), a key gene involved in B cell activation and maturation (15). In addition to this initial activation module, we identified a secondary activation state comprising several poorly understood TFs, including BHLHE40, CEBPE/Z, ZBTB33, and ZHX1 (Module 3). We also identified dynamic expression and chromatin activity in GC B cells, including one module that decreases through GC exit and plasma cell differentiation (Module 4; HNF1B, EBF1, SMAD2, POU2F1, MEF2B) and one module that is maintained or increases during commitment to the plasma fate (Module 5; NR2F6, FOXO4, JDP2, MSC). In contrast, a transcriptional regulatory module containing master plasma cell regulators such as IRF4, PRDM1 and XBP1 (Module 6) exhibited reduced accessibility at target sites within GC B cells compared to both naive and plasma populations, suggesting that these sites may be actively repressed to prevent inappropriate or premature commitment to the plasma fate during affinity maturation in the GC.

### Integration of secondary lymphoid organ datasets with bone marrow and peripheral blood single-cell transcriptome and epigenome atlases

Other scRNA-seq analyses have recently demonstrated that tonsils are a transferable model tissue to study secondary lymphoid organs and adaptive immune responses more generally (10, 11, 16). In contrast to circulating or bone-marrow resident lymphocyte populations, immune cells within secondary lymphoid organs exist in a range of activation and maturation states, including GC-associated populations, that may reflect varied tissue niches, cell-cell communication and cytokine signaling. To examine the potential relevance of tissue-specific gene expression and chromatin-based regulatory activities, we integrated our tonsillar scRNA-seq and scATAC-seq datasets with those from publicly available bone marrow and peripheral blood immune cell atlases (17) to generate an overview of immunopoiesis comprising data for 60,639 and 91,510 high quality cells for scRNA-seq and scATAC-seq respectively (Fig3a, S5, TableS7-8). As expected, activated B cells, GC-associated lymphocytes (GC B and Tfh cells) and tissue-resident macrophages were strongly enriched in secondary lymphoid organs, while progenitor populations and circulating monocytes were enriched in the bone marrow and peripheral blood (Fig3b). In addition to differences in the frequency of immune cell subsets, we also examined if there might be differences between circulating or tissue resident B cells. We found significant differences in both the chromatin accessibility and gene expression of naive and memory B cells in the tonsil compared to matched populations in the periphery (Fig3c-d, S6). In particular, chromatin accessibility profiles of tonsillar B cells were enriched with POU2F2 (also known as OCT2) motif sequences (Fig3e), a TF known to be important in the regulation of humoral B cell responses (18). These tissue-specific phenotypes likely reflect differences in cytokine exposure and microenvironment of the tonsil compared to circulating blood and highlight that it is essential to examine immune cell populations across varied tissue contexts, even for a single cell type.

**Figure 3.**
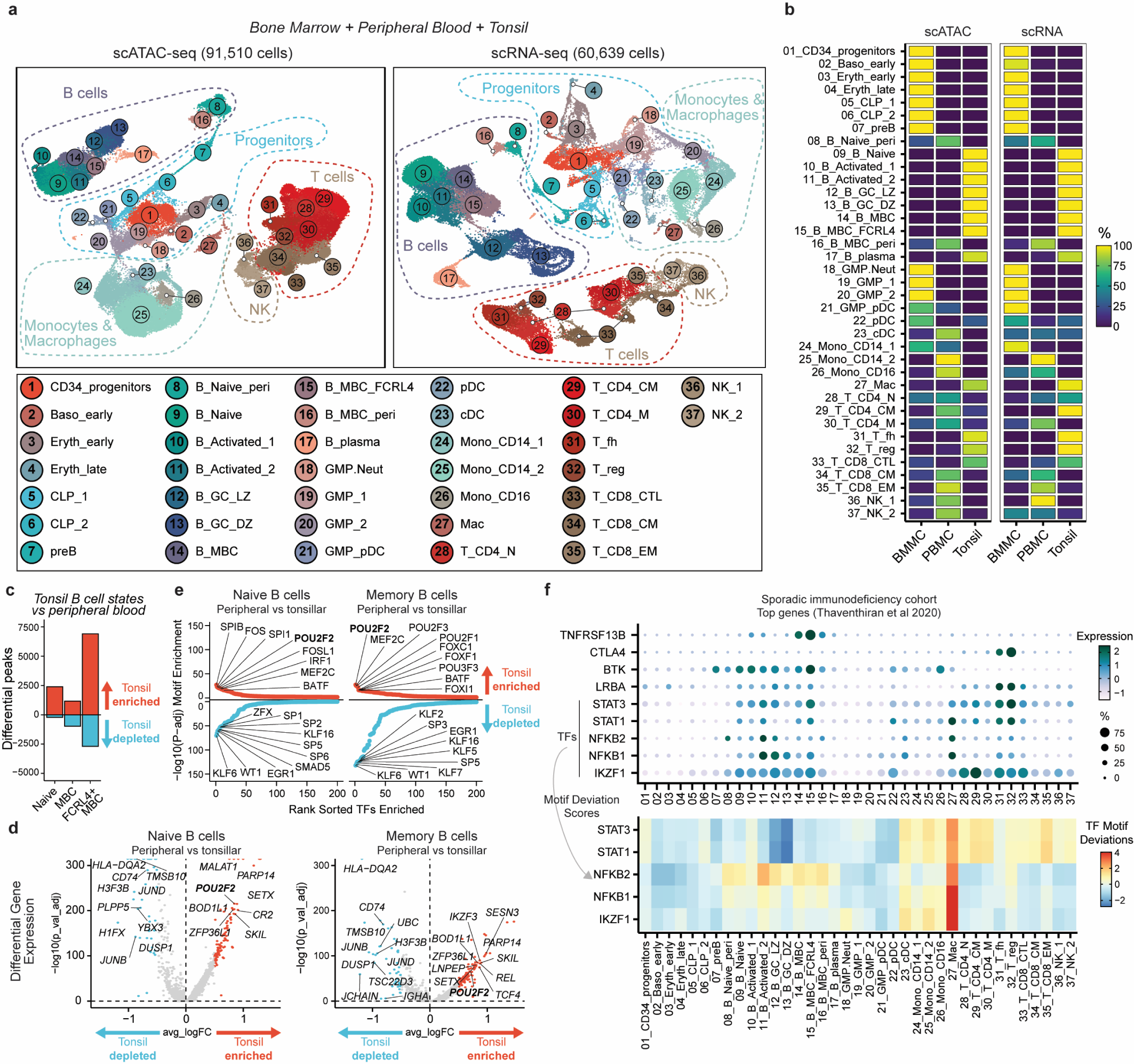
Integrated single-cell transcriptomics and epigenomics of human bone marrow, peripheral blood and tonsillar immune cell states. a) UMAP of integrated scATAC-seq and scRNA-seq for human bone marrow, peripheral blood and tonsils. b) Relative frequency of cell type clusters in A) across different tissues. c) Differential scATAC-seq peak analysis of tonsillar compared to peripheral blood/bone marrow-enriched naive and memory B cell clusters. FCRL4+ MBC cluster was compared to peripheral blood-enriched MBC cluster. d) Differential gene expression analysis of tonsillar compared to peripheral blood/bone marrow-enriched naive and MBC clusters in integrated scRNA-seq dataset. Selected genes are annotated. e) Ranking of TF motif deviation enrichment within tissue-enriched (red, upper) or tissue-depleted (blue, lower) peaks naive and memory B cells. f) Expression of top genes identified to be mutated by whole genome sequencing in a sporadic immunodeficiency cohort (19). For TFs, motif deviation scores are also provided.

Finally, we examined the cell type-specific expression of nine genes recently identified to be most commonly mutated within a sporadic primary immunodeficiency cohort (Fig3f) (19). Two genes, TNFRSF13B and CTLA4 were relatively cell type-specific in their expression pattern. TNFRSF13B (encoding TACI) was most highly expressed in memory B cells, particularly tonsillar FCRL4^+^ memory B cells. Patients with TNFRSF13B mutations have fewer memory B cells expressing class-switched antibodies (20). CTLA4 expression peaked in Tfh and Treg populations as expected. In contrast, BTK, LRBA, and the TF genes STAT1, STAT3, NFKB1, NFKB2 and IZKF1 were broadly expressed across varied subsets. We used our scATAC-seq data to examine the enrichment of their motif sequences in accessible chromatin to determine which cell type might be most sensitive to altered activity of these TF genes. This revealed that tonsillar myeloid cells (labelled here primarily as macrophages) had the highest activity of these immunodeficiency-associated TFs (Fig3f), although we observed enrichment of NFKB2 in activated B cells (Fig2f, 3f) and STAT1/STAT3 in circulating monocyte and T cells (Fig3f).

### Identification of fine-mapped autoimmune GWAS variants in cell type-specific chromatin

Our integrated scRNA-seq and scATAC-seq atlas of immune cell populations in bone marrow, peripheral blood and tonsils provided a unique opportunity to understand the regulatory potential and cell type-specificity of autoimmune-associated genetic variants across a broad diversity of immune cell types. By examining 12,902 fine-mapped SNPs, of which 9,493 were significantly associated with disorders of the immune system (1, 21), we found that our single-cell accessibility profiles of immune cells were broadly enriched in immune-related genetic variants compared to non-immune related traits and background genetic variation (Fig4a, S7a-b). Importantly, we found specific enrichment of disease-specific genetic variants in different immune cell lineages or subsets (Fig4b, S7c-d). For example, we found a strong enrichment of genetic variants associated with Kawasaki disease and lupus in chromatin accessibility maps of the B cell lineage, particularly tonsillar naive and memory B cells, as well as enrichment of genetic variants associated with alopecia, autoimmune thyroiditis, systemic sclerosis and Behçets disease in cytotoxic lymphocyte regulatory elements (Fig4b, S7c-d). In contrast, genetic variants associated with multiple sclerosis were enriched in both B and T cell-specific chromatin, perhaps reflecting the multigenic nature and complex etiology of this disease (Fig4b, S7c-d).

**Figure 4.**
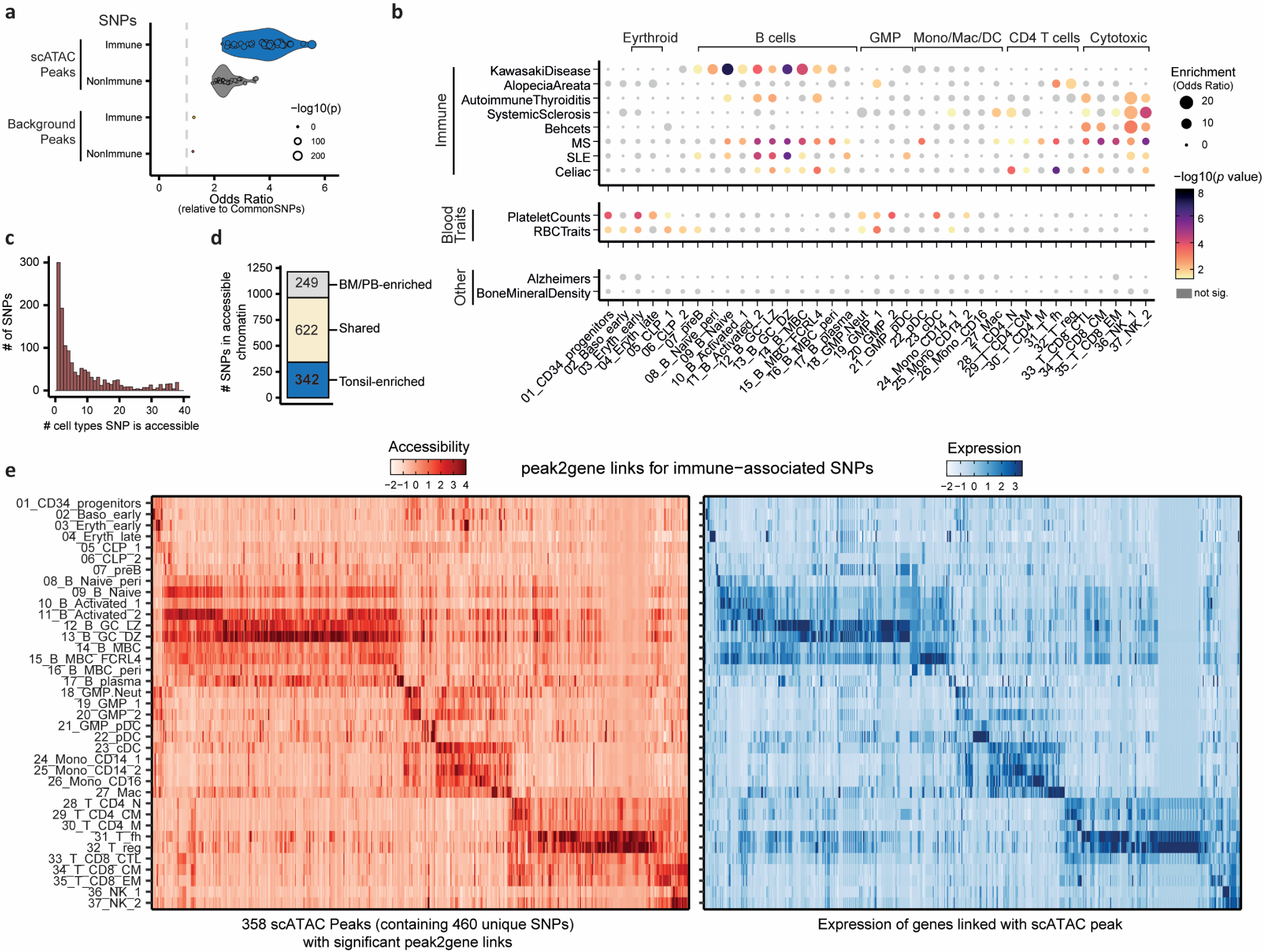
Autoimmune-associated genetic variants enriched in immune cell chromatin accessibility maps. a) Fisher enrichment test of immune-associated fine mapped genetic variants, compared to common genetic variants, for chromatin accessibility scATAC peaks across 37 immune cell populations. Results for non-immune traits and background control peaks are shown. Dot size conveys significance (-log10(p value)). b) Fisher enrichment test for trait-specific SNPs, compared to the complete fine-mapped SNP set, within cell type-specific chromatin accessibility peaks. Dot size conveys enrichment (odds ratio) and color denotes significance of enrichment. c) Frequency histogram of immune-associated SNPs that fall within chromatin accessibility peaks across 37 immune cell types. d) Tissue-specificity of chromatin accessibility peaks overlapping autoimmune SNPs. e) Chromatin accessibility of peaks containing >1 immune-associated SNP (scATAC; left) for which at least one significant peak2gene linkage is identified. Expression of linked genes (scRNA; right) is also plotted. Accessibility or expression counts are scaled by peak or gene respectively.

Of the 1213 immune-related SNPs that overlapped with accessible chromatin peaks in our atlas (TableS9), many were localized in cell type-or lineage-specific chromatin (Fig4c). Importantly, 342 (28.2%) of these SNPs fell within accessible chromatin only identified in tonsil-enriched immune subsets (Fig4d), demonstrating the value of our tonsillar immune cell atlas for interpretation of GWAS genetic variants. We next predicted the putative gene targets of these genetic variants by using our integrated scRNA-seq and scATAC-seq to identify highly correlated accessibility at chromatin regions to nearby gene expression (12, 17). This enabled us to examine 358 chromatin accessible regions (containing 460 unique immune-linked SNPs) for which we could identify significant peak-to-gene linkages (Fig4e). These linkages revealed cell type-specific and shared patterns of both the chromatin accessibility of autoimmune genetic variants and expression of putative gene targets, providing a powerful resource to explore the potential regulatory mechanisms of these genetic variants and their contribution to autoimmune disease (Fig5-6).

**Figure 5.**
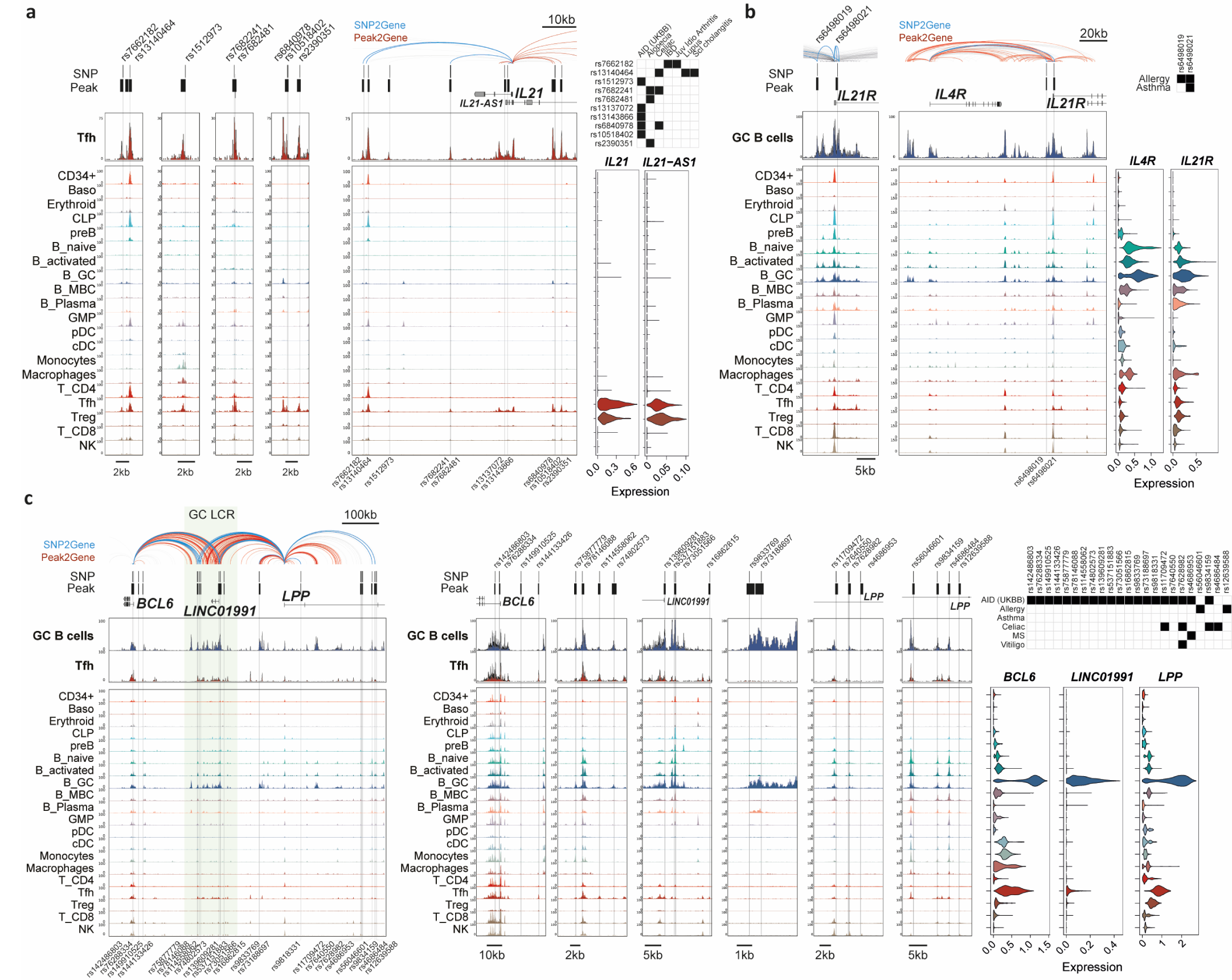
Chromatin regulatory landscapes of GC-specific autoimmune risk variants. a) Genomic snapshot of fine-mapped autoimmune-associated GWAS variants that localize to accessible chromatin in the integrated human bone marrow, peripheral blood and tonsil scATAC-seq atlas. High resolution of individual SNP loci and larger view of the IL21 locus are shown, with significant peak2gene linkages colored in red and significant links between SNPs and gene promoters (SNP2gene) in blue and bold. Significant associations between individual SNPs and autoimmune diseases are shown in black boxes and gene expression is shown as violin plots for matched populations in the scATAC tracks. AID; autoimmune disease. IBD; inflammatory bowel disease. Juv Idio Arthritis; Juvenile idiopathic arthritis. Scl cholangitis; Primary sclerosing cholangitis. b) Same as A), at the IL4R/IL21R locus. c) Same as A), at the BCL6/LPP locus. A germinal center (GC)-specific locus control region (LCR) is highlighted in green. MS; multiple sclerosis

### Chromatin regulatory activity at immune-associated genetic variants predicts importance of GC activity in autoimmunity

Many studies examining the relationship between immune-associated genetic variants and their regulatory activity with functional genomics methods such as ATAC-seq or ChIP-seq have been limited to studying peripheral immune cell populations. This limitation is likely significant, given our knowledge that many lymphocyte maturation and antibody-based selection events occur in secondary lymphoid organs, and that GC-derived autoantibody production is a feature of many autoimmune diseases. Although we found examples of genetic variants in cell type-specific chromatin across diverse immune subsets (Fig4e, S8-S9; e.g. GZMB/GZMH, NKX2-3, COTL1/KLHL36, KSR1/LGALS9, TNFRSF1A/LTBR), we observed a striking enrichment of fine-mapped autoimmune variants in chromatin accessibility regions specific to GC-associated B and T populations, such as GC B cells and Tfh cells (Fig4e), including the IL21, IL21R/IL4R, BCL6/LPP, CD80, PRAG1, SLC38A9, VAV3/SLC25A24, DLEU1/DLEU1/TRIM13 loci (Fig5, 6, S10-S11).

**Figure 6.**
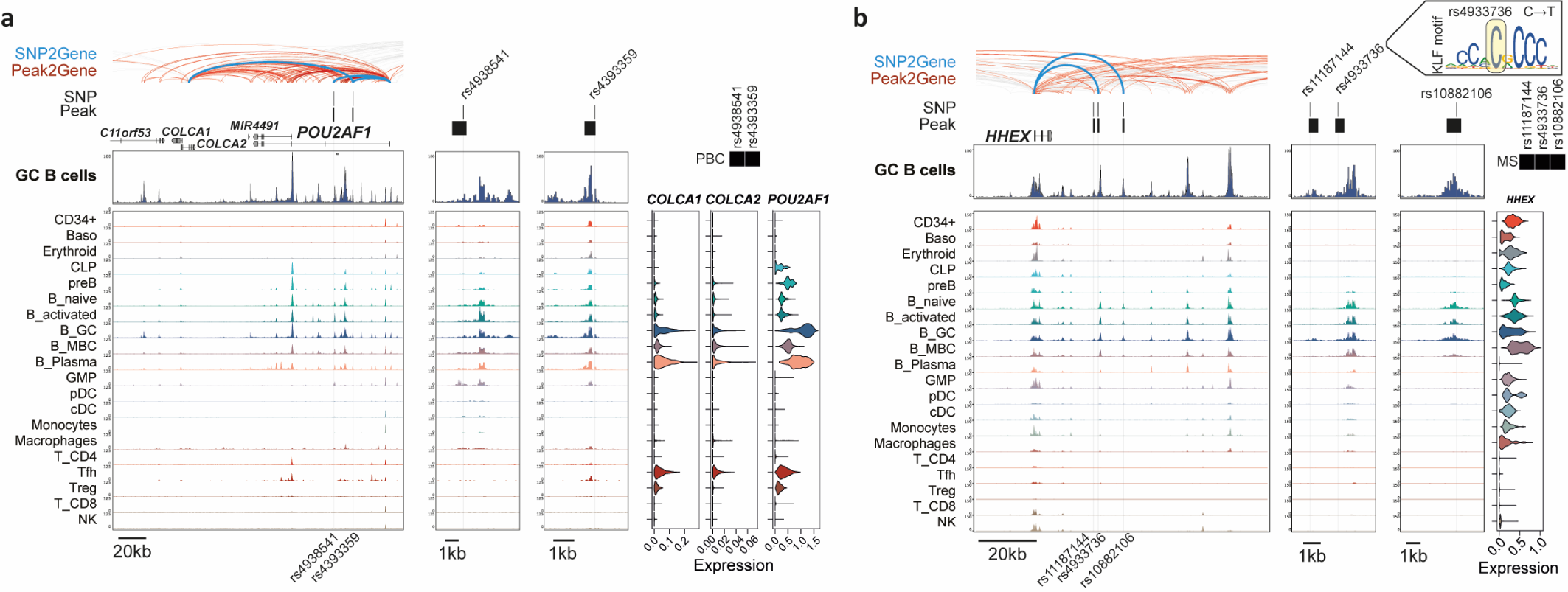
Autoimmune risk variants at transcription regulator genes POU2AF1 and HHEX. a) Genomic snapshot of fine-mapped autoimmune-associated GWAS variants at the POU2AF1 locus that localize to accessible chromatin in the integrated human bone marrow, peripheral blood and tonsil scATAC-seq atlas. Significant peak2gene linkages colored in red and significant links between SNPs and gene promoters (SNP2gene) in blue and bold. Significant associations between individual SNPs and autoimmune diseases are shown in black boxes and gene expression is shown as violin plots for matched populations in scATAC tracks. PBC; primary biliary cirrhosis b) Same as A), at the HHEX locus. MS; multiple sclerosis

We identified GC-specific regulatory elements at the IL21 locus and the locus of its receptor IL21R (Fig5a-b, FigS1). Cytokine signaling by IL-21, primarily secreted by Tfh cells, is essential for B cells to form and participate in normal GC reactions. B cells respond to IL-21 through the IL-21 receptor (IL-21R). We identified several fine-mapped SNPs at the IL21 locus highly correlated with both chromatin accessibility and gene expression at the IL21 promoter (Fig5a). These SNPs exhibited Tfh-specific chromatin accessibility, although one SNP, rs13140464, was also highly accessible in several progenitor populations. These fine-mapped SNPs at IL21 have been associated with alopecia (1), juvenile idiopathic arthritis or autoimmunity more generally (21), and some of these same SNPs are also significantly associated with celiac disease (rs7682241, rs6840978) (22), inflammatory bowel disease (rs7662182) (23), alopecia (rs2390351) (24), primary sclerosing cholangitis (rs13140464) (25) and lupus (rs13140464) (26). Conversely, we also found two fine-mapped SNPs in strong linkage disequilibrium (rs6498021, rs6498019) located in close proximity to IL21R in B cell-specific chromatin accessibility regions that have been linked with allergy (1) and/or asthma (27) (Fig5b, S12). As well as predicted interactions with IL21R, these two SNPs were also predicted to interact with nearby IL4R gene, encoding the IL-4 receptor (IL-4R), which, similar to IL21R, was most highly expressed in GC B cells and is vital for T cell-dependent maturation of B cells.

### Autoimmune risk variants within a GC-specific locus control region

Our analysis of genetic variants linked with autoimmunity identified a concentration of recently fine-mapped autoimmune-associated SNPs from the UKBB databank (21) in a GC-specific locus control region (LCR) (28) located between BCL6 and LPP (Fig5c, S12). Of the genetic variants that fell within accessible chromatin across this locus, there were associations with celiac disease (rs11709472 (29), rs7628982 (UKBB), rs9834159 (30), rs4686484 (1)), allergy (rs56046601 and rs12639588 (1)), multiple sclerosis (rs4686953 (formerly rs66756607) (31, 32)), asthma (rs7640550 and rs7628982 (33)) and vitiligo (rs7628982 (34)). Many of these SNPs were present in chromatin accessible regions specific to GC B or Tfh cells, in which BCL6, LPP and the long non-coding RNA at the LCR (LINC01991) are most highly expressed (Fig5c). We report significant correlations in chromatin accessibility between many of these SNPs (and the LCR in general) for both BCL6 and LPP, consistent with chromosome conformation interactions detected in GC B cells between this LCR and the BCL6 promoter (28, 35). Importantly, deletion of this LCR has been shown in mouse models to lead to defects in GC B cell formation (28), presumably through its transcriptional regulation of BCL6, one of the master regulatory TFs required for both GC B cells and Tfh cells. These observations suggest that association of this locus with autoimmunity is primarily driven through GC B and Tfh defects. However, some genetic variants (rs142486803, rs76288334, rs78146088) were accessible across many different immune lineages, as well as rs4686484 that was previously proposed to be located in a B cell-specific enhancer (30), revealing an additional layer of complexity to this autoimmune regulatory locus.

### Autoimmune risk variants at the loci of transcriptional regulators POU2AF1 and HHEX

We identified cell type-specific chromatin accessibility at autoimmune risk variants across loci for many regulatory TFs or transcriptional regulators including POU2AF1, HHEX, ETS1, STAT4, IKZF3, NKX2-3 and IRF8 (Fig6, S8, S13-14), in addition to the GC master regulator BCL6 (Fig5c). Of particular interest were POU2AF1 and HHEX, which have recently been proposed to control memory B cell fate selection in the GC (36, 37). POU2AF1, also known as OCT binding factor 1 (OBF1), is a largely B cell-specific transcriptional coactivator with no intrinsic DNA binding activity that interacts with TFs POU2F1 (OCT1) and POU2F2 (OCT2). It is indispensable for formation of GCs and GC-dependent B cell maturation (38–40). We found two genetic variants associated with primary biliary cirrhosis/cholangitis (PBC) (rs4938541 and rs4393359 (1, 41)) within B cell-specific accessible chromatin and observed that POU2AF1 expression peaks in GC B cells (42) (Fig6a). Our analysis of B cell activation dynamics predicted POU2F1/POU2F2 as regulators in GC B cells (Fig2) and POU2F2 is more highly expressed in tonsillar B cells compared to those circulating in peripheral blood (Fig3), suggesting that B cells within lymphoid tissues are likely to be most sensitive to altered POU2AF1 levels.

HHEX has recently been reported to be an essential regulator of the memory B cell fate decision by GC B cells (37), although its potential involvement in autoimmune disease is not known. Our integrated epigenomic and transcriptomic analyses identified three fine-mapped SNPs at the HHEX locus that fell within B cell specific-accessible chromatin, were predicted to regulate HHEX through peak-to-gene correlation analysis and were associated with multiple sclerosis (MS) (rs11187144, rs4933736, rs10882106 (1)) (Fig6b). We also identified linkages between these SNPs and neighboring genes KIF11 and EXOC6 (FigS15). We note that rs4933736 is predicted to fall within a predicted KLF TF binding site (Fig6b), providing a potential mechanism for disruption of HHEX expression.

## Discussion

Here, we generate paired transcriptome and epigenome atlases of immune cell subsets in the human tonsil, a model system to study the GC reaction which is a major site for peripheral tolerance to prevent autoimmunity, and adaptive immunity to respond to infection. We define gene expression and gene regulatory elements across dynamic immune cell states and examine the regulatory potential of transcription factors in these populations. Our analyses highlight the importance of examining cellular populations in secondary lymphoid tissues, compared to peripheral blood or bone marrow of previous studies, to understand how regulatory activity at non-coding genetic variants in crucial, yet transient, populations might contribute to autoimmune disease.

Our single-cell transcriptomic analysis identified a rare B cell population that expresses high levels of IFN-induced gene expression (Fig1). Unfortunately, we were unable to identify this rare B cell population in our scATAC profiling to explore how it may be linked to different autoimmune traits at the chromatin level. One of the genes most highly expressed by the IFN-responsive B cells was IFI44L. Splice and missense genetic variants at the IFI44L locus (rs1333973 and rs273259) have previously been linked with neutralizing antibody titers to the measles vaccine (43), and type I interferon-positive B cells have previously been implicated in the development of autoreactive B cells (44). These observations suggest this rare and poorly characterized B cell state may be involved in B cell-mediated antibody responses to vaccines and/or the development of autoimmunity.

The joint analysis of gene expression with chromatin accessibility landscapes allowed us to predict putative TF regulators in both steady state and dynamic immune cell populations, including temporally dynamic TFs during B cell activation and GC formation. We identified a secondary B cell activation state, after an initial NFκB-associated activation presumably linked with strong BCR activation and/or T cell help (45). One particularly interesting candidate for future study will be BHLHE40, which has previously been shown to be required for the transition from an activated state prior to entry into the GC (46–48) and is capable of binding key regulatory elements at the immunoglobulin heavy chain locus (10). How this, and other putative regulators we identify in this secondary activation state (such as CEBPE/Z, ZBTB33, and ZHX1) may contribute to the transition from the activated B cell state to a GC-associated gene expression program will be an important question for future mechanistic studies.

The molecular mechanisms by which many GWAS-identified genetic polymorphisms contribute to autoimmune disease remain poorly understood. Here, we have examined diverse immune cell lineages to reveal that many autoimmune disease-associated genetic variants are localized within chromatin most accessible in GC B and T cell populations, including genes important for B cell activation (CD83, CD80, IL21), survival and participation in the GC (IL21R, IL4R, BCL6) and fate selection (POU2AF1, HHEX, IRF8). While our findings do not exclude regulatory potential at these or other autoimmune-associated loci in stromal cell populations (49), they strongly implicate lymphocyte-intrinsic dysfunctional GC responses as a major feature in the etiology of autoimmune disease.

While at some loci our analyses predict disruption of a TF binding site by genetic variants (HHEX, CD83), our current analyses are not able to predict whether specific polymorphisms might positively or negatively regulate gene expression at their nearby targets. Current eQTL databases have relied upon whole tissue expression data, typically from adult (i.e. non-pediatric) lymphoid tissue like spleen or peripheral blood which lack adequate representation of GC cellular populations. Single-cell multiomics and eQTL analyses in varied healthy and diseased immune organs and model systems will be essential to provide further mechanistic insights (16, 50). Finally, new advances in neural-network derived methods may also prove useful to quantitatively predict effects on gene expression in cell type-resolved chromatin accessibility maps (14, 51).

Dysregulated GC responses are well known to contribute to the development of autoimmune disease (52). For example, defects in GC formation or activity would provide an opportunity for the expansion of self-reactive B cells that are normally inhibited in the periphery of healthy individuals (53). In other contexts, B cell activation has been linked with the induction of autoimmunity in animal models (54). The importance of normal GC function in preventing autoimmunity is also illustrated in an experimental mouse model of lupus where the B cell-specific depletion of IL-21R, essential for normal GC responses, prevents the development of autoantibodies and disease (55). Although the translation of mouse models to human autoimmune diseases can be challenging, we were able to identify several genetic variants in Tfh-or GC B cell-enriched gene regulatory elements at both IL21 and IL21R that were associated with varied immune disease traits (Fig5).

While defective GC responses may contribute to autoimmunity in one manner, over-activation of the GC response may act in another. The development of transient GC-like lymphoid follicles in non-lymphoid tissue (termed ectopic GCs) has been associated with site-specific inflammation in autoimmune diseases and may contribute to loss of tolerance by promoting maturation of self-reactive B cell clones (56). Analysis of B cells from ectopic GCs in several autoimmune diseases provide evidence of site-specific clonal expansion and somatic hypermutation of antibody genes, and an absence of normal GC regulation (57–59). Single cell analyses of “defective” and “ectopic” immune structures in different autoimmune diseases will be essential to understand how the regulatory and gene expression dysfunction we predict in the normal immune cell landscape may drive autoimmunity through altered GC response dynamics.

## Materials and Methods

### Human ethics, tissue collection and preparation

Tonsil samples were collected from children and adults undergoing routine tonsillectomy. All participants provided written informed consent and the protocols were approved by Stanford University’s Institutional Review Board (protocol numbers 30837 and 47690). Whole tonsils were collected in saline and processed within four hours of receipt. Tissues were treated with penicillin, streptomycin, and normocin for 30 minutes on ice and heavily clotted or cauterized areas of the tissue were removed. Tonsils were then dissected into small pieces (roughly 5-8 pieces per tonsil) before mechanical dissociation through a 100 µm cell strainer using a syringe plunger. Mononuclear cells were isolated by ficoll density gradient centrifugation (GE Healthcare) and the buffy coats were collected. Cells were cryopreserved in 90% fetal bovine serum, 10% DMSO until use. Four additional cryopreserved tonsil samples at Queen Mary University of London included for scATAC-seq analyses were prepared as described previously (10) under approval from North West/Greater Manchester East Research Ethics Committee (17/NW/0664).

### CyTOF staining and analysis

Cryopreserved samples were thawed in pre-warmed cell culture medium (RPMI1640 with 10 % FBS, non-essential amino acids, sodium pyruvate, antibiotics), washed, and rested for 1 hour at 37°C in culture medium supplemented with DNase (25 U/ml). Cells were then washed and resuspended in FACS buffer (PBS with 0.1% w/v bovine serum albumin, 2 mM EDTA, 0.05% v/v sodium azide). Individual donor samples were barcoded using a combination of metal-tagged CD45 antibodies, combined into barcoded pools, stained for surface antibody markers, and treated with cisplatin for viability staining as described (60). Samples were then fixed overnight with 2% paraformaldehyde diluted in PBS. The next day, cells were permeabilized using a permeabilization buffer (eBioscience), stained with a DNA intercalator for 30 minutes, and washed. Just prior to CyTOF data collection, samples were washed three times with PBS, then three times with MilliQ water. Barcoded pools were run on a CyTOF2 instrument (Fluidigm) and fcs files were exported for analysis in FlowJo software. Live intact singlets were gated and samples were manually debarcoded using combinations of CD45 channels (5-choose-2 scheme) and individual donor samples were exported as separate fcs files before dimensionality reduction analyses.

### Single-cell library preparation, sequencing and alignment

Tonsillar immune cells were loaded on to the 10X Genomics Chromium according to the manufacturer’s protocol using either the single-cell 3’ kit (v3) or the single-cell ATAC kit (v1). Library preparation for both assays was performed according to the manufacturer’s protocol prior to sequencing on either the Illumina NovaSeq 6000 or NextSeq 500 platforms. scRNA-seq libraries were sequenced with 28/10/10/90 bp cycles while scATAC-seq libraries were sequenced with 70/8/16/70 bp read configurations. BaseCall files were used to generate FASTQ files with either cellranger mkfastq (v3; 10X Genomics) or cellranger-atac (v1; 10X Genomics) prior to running cellranger count with the cellranger-GRCh38-3.0.0 reference or cellranger-atac count with the cellranger-atac-GRCh38-1.1.0 reference for scRNA-seq and scATAC-seq libraries respectively.

### Quality control, integration and cell type annotation of tonsillar scRNA-seq

Gene expression count matrices from cellranger were processed with Seurat (v3.0.2) (61, 62) for genes detected in greater than 3 cells. Cell barcodes were filtered based on the number of genes per cell (between 200-7500), percentage of mitochondrial reads per cell (0-20 %) and the number of ADTs (less than 4000). Initial data quality control was performed separately on data from each biological sample. Data from technical replicate libraries were combined, normalized with SCTransform (63) before highly variable gene identification and PCA dimensionality reduction. Jackstraw plots were visually assessed to determine the number of principal components (PCs) for subsequent analysis: Tonsil1 = 11, Tonsil2 = 13, Tonsil3 = 12. Preliminary clusters were identified (FindClusters; res = 0.8) before computing UMAP dimensionality reduction and identifying putative doublets with DoubletFinder (64) (sct=TRUE, expected_doublets=3.9%). Pre-processed Seurat objects were then merged, with SCTransform normalization and PCA computation repeated using all variable features (except for IGKC, IGLC, IGLV, HLA, and IGH genes). Batch correction was performed with harmony (65). UMAP dimensionality reduction and cluster identification were performed (27 PCs, res = 0.8). For higher resolution analysis of B cells and T cells, data from B or T cells only were processed separately, with repeated variable gene identification (removing IGKC, IGLC, IGLV, HLA, and IGH) before repeated PCA, batch correction with Harmony, UMAP reduction and cluster identification (30 PCs, res = 0.6 for B cells; 20 PCs, res = 0.6 for T cells). Gene expression markers for clusters were identified (FindAllMarkers; log fold change > 1, adjusted p value < 0.05). Imputation of gene expression counts (for plotting only) was performed with MAGIC (66).

### scATAC-seq quality control, batch correction and integration with scRNA-seq datasets

Mapped Tn5 insertion sites (fragments.tsv files) from cellranger were read into the ArchR (v0.9.4) R package (12) retaining cell barcodes with at least 1000 fragments per cell and a TSS enrichment score > 4. Doublets were identified and filtered (addDoubletScores and filterDoublets, filter ratio = 1.4) before iterative LSI dimensionality reduction was computed (iterations = 2, res = 0.2, variable features = 25000, dim = 30). Sample batch correction was performed with harmony (65). Clustering was then performed on the harmony corrected data (addClusters, res = 0.8) before UMAP dimensionality reduction (nNeighbors = 30, metric = cosine, minDist = 0.4). One cluster enriched for high doublet scores (cluster 7) was removed. Tonsillar scRNA-seq gene expression and cell type identities were integrated with the tonsillar scATAC data using harmony-corrected data with ArchR as previously described (12). A preliminary cell type annotation was performed using gene accessibility scores of known cell type markers. To improve cell type assignment of closely related cell types, we performed a constrained integration, grouping GC B cell clusters, other B cell clusters and non-B cell clusters together during addGeneIntegrationMatrix. The most common predicted cell type from the integration with RNA expression in each previously identified ATAC-seq cluster was used to annotate scATAC cluster identity. The quality of mapping between the RNA and ATAC was confirmed by identifying marker gene scores in scATAC clusters using getMarkerFeatures. Additionally, cluster annotations derived from scATAC-only analysis were compared with cluster annotations derived from scRNA-seq integration.

For high resolution clustering of B and T cell subsets (Fig2), scATAC clusters identified as B cells or T cells following scATAC/scRNA integration were subset, and use to recompute iterative LSI dimensionality reduction as described above, except 30 dimensions were used for B cell analysis. Batch correction, cluster identification and UMAP reduction were also performed as above, except that minDist = 0.1 (T cells) or 0.3 (B cells). Integration of B cell and T cell scATAC-seq datasets with gene expression and high resolution cluster annotations was performed using the T cell-or B cell-specific scRNA-seq Seurat objects as previously described with addGeneIntegrationMatrix in ArchR. Integration between assays were constrained with the following broad groups: B cell subgroups; plasmablasts, memory, naïve/activated and GC B cell clusters, T cells; CD8+/cytotoxic T cells and remaining T cell clusters.

### Peak calling and inference of transcription factor activity in scATAC-seq datasets

Single-cell chromatin accessibility data were used to generate pseudobulk group coverages based on high resolution cluster identities of scATAC-seq datasets before peak calling with macs2 (67) using the addReproduciblePeakSet in ArchR. A background peak set controlling for total accessibility and GC-content was generated using addBgdPeaks. Chromvar (68) was run with addDeviationsMatrix using the cisbp motif set to calculate enrichment of chromatin accessibility at different TF motif sequences in single cells. To identify correlation between the gene expression and transcription factor activity, RNA-expression projected into the ATAC subspace (GeneIntegrationMatrix) and the Chromvar deviations (MotifMatix) were correlated using correlateMatrices. A correlation of greater than 0.25 was used to determine if the TF correlated between expression and activity and the list of correlated TFs was further subset by only including TFs that were expressed in at least 25 percent of cells in one or more cell type cluster. To analyze transcription factor activity during B cell activation, GC entry and plasma differentiation, the harmony-corrected B cell ArchR object was subjected to “addTrajectory” from ArchR using the following user-defined trajectory as a guide: Naive→Activated→LZ GC→DZ GC→Plasmablasts. Gene expression and Chromvar deviation scores were correlated throughout pseudotime using correlateTrajectories (corCutOff = 0.25, varCutOff1 = 0.25, varCutOff2 = 0.25) and visualized using plotTrajectoryHeatmap. “Peak-to-gene links” were calculated using correlations between peak accessibility and integrated scRNA-seq expression data using addPeak2GeneLinks.

### Integration of tonsil scATAC-seq and scRNA-seq with bone marrow and peripheral blood datasets

Published bone marrow and peripheral blood scRNA-seq and scATAC-seq (17) were aligned to the hg38 genome as described above. Additional hg38-aligned PBMC scATAC-seq datasets were downloaded from 10X Genomics (https://support.10xgenomics.com/single-cell-atac/datasets).

#### scRNA-seq

Cellranger gene expression matrices were used to sum and quantify mitochondrial gene expression before mitochondrial genes were removed from the gene expression matrices. Similarly, V, D and J gene counts from T cell and immunoglobulin receptors were summed and removed from matrices. Closely related IgH constant region genes were also summed and removed (IgG1-4, IgA1-2). Cell barcodes expressing >200 genes and genes detected in >3 cells were then processed in Seurat (61, 62), with doublet prediction using default settings with scrublet (69) (expected doublet frequency 8x10^-6^ X 1000 cells). Predicted doublets were removed, and cell barcodes with <750 or >30000 UMIs, <500 or >6000 genes detected, or >20% mitochondrial gene expression were also removed. Individual datasets were then merged together, before normalization and batch correction with SCTransform (3000 variable features) and scoring of cell cycle phase with Seurat. “IGLsum”, “IGKsum”, “IGHG”, “IGHA”, “IGHM”, and “IGHD” were subsequently removed from highly variable gene list so they would not contribute to downstream dimensionality reductions. PCA was then computed before UMAP reduction (n.neighbors = 20, min.dist = 0.35, dims = 1:50), nearest neighbor identification (FindNeighbours; dims = 1:50) and cluster identification (FindClusters; res = 1.75). Some additional subclustering was performed to better match cell type annotations from previous tonsil analysis (this study) and peripheral blood/bone marrow analysis (17). In general, previous annotations were closely adhered to and confirmed by examination of known cell type-specific gene expression markers. Differential gene expression between clusters was performed with FindAllMarkers or FindMarkers, with padj < 0.05 and avg_logFC > 0.5. Imputation of gene expression counts (for plotting only) was performed with MAGIC (66).

#### scATAC-seq

Cellranger-derived fragments.tsv files of tonsil, peripheral blood and bone marrow samples were processed with ArchR (12) (createArrowFiles; filterTSS = 6, filterFrags = 1000, minFrags = 500, maxFrags = 1e+05). Doublets were identified (addDoubletScores; k=10) and removed with a filterRatio = 1.4, before additional filtering of cell barcodes to remove those with TSSEnrichment < 6, < 10^3.25^ or > 10^5^ fragments per barcode, nucleosome ratio of > 2.5, ReadsInBlacklist > 800, or BlacklistRatio > 0.009. Preliminary LSI reduction was performed with addIterativeLSI (corCutOff = 0.25, varFeatures = 30000, dimsToUse = 1:40, selectionMethod = “var”, LSIMethod = 1, iterations = 6, filterBias = FALSE, clusterParams = list(resolution = c(0.1,0.2,0.4,0.6,0.8,1), sampleCells = 10000, n.start = 10). To account for differences in sequencing coverage, Harmony batch correction (corCutOff = 0.25, lambda = 0.75, sigma = 0.2) was performed using library ID for tonsil samples, public 10X Genomics PBMC datasets and sample BMMC_D6T1, while remaining samples from Granja et al were treated as a single batch. Preliminary identification of clusters (addClusters; res = 1.5) identified two poor quality clusters enriched with doublets (C38, C7). These were removed from subsequent analysis. Quality controlled datasets were then subjected to new LSI dimensionality reduction and Harmony batch correction with the same settings, before computing UMAP (RunUMAP; nNeighbors = 80, minDist = 0.45, seed = 1) and identifying cell type clusters with at least 80 cells (addClusters (method = “Seurat”, res = 1.1 or 1.5, nOutlier = 80). Broad lineages were first annotated to help with integration and transfer of scRNA expression. Normalized, non-corrected scRNA expression counts and annotated cell types were transferred to nearest neighbour scATAC cells using addGeneIntegrationMatrix (sampleCellsATAC = 10000, nGenes (RNA) = 4000, sampleCellsRNA = 10000) with a constrained integration to the following groups: CD4T_cells, CD8T_cells, GC_PB, MBC_B_cells, Myeloid_cells, NaiveAct_B_cells, NK, Peripheral_B_cells, Progenitors. Accessibility gene scores and transferred RNA expression counts were imputed with addImputeWeights(corCutOff = 0.25). Cell type clusters were carefully annotated with a combination of pre-existing annotations from Granja and tonsil immune cell scATAC data (this study), transferred cell annotations from scRNA-seq and examination of known subset markers.

Pseudobulk group coverages of cell type clusters were calculated with addGroupCoverages and used for peak calling using macs2 (addReproduciblePeakSet in ArchR). A background peak set controlling for total accessibility and GC-content was generated using addBgdPeaks. Cell type-specific marker peaks were identified with getMarkerFeatures with the wilcoxon test and controlled for TSSEnrichment and fragment count. Peak accessibility was deemed significantly different between clusters if FDR < 0.05 and log2fc > 0.56. Motif annotations and enrichment were calculated as described above with addMotifAnnotations and addDeviationsMatrix. “Peak-to-gene links” were calculated using correlations between peak accessibility and integrated scRNA-seq expression data using addPeak2GeneLinks (corCutOff = 0.25).

### Analysis of fine-mapped GWAS variants

The results of two independent GWAS fine-mapping studies (1, 21) (https://www.finucanelab.org/data) were combined. PICS SNPs from both immune and non-immune traits were included in analyses, while only SNPs from the study mapping the UKBB resource that were associated with a combined autoimmune disease trait (AID; labelled as AID_UKBB) were included. This provided a total of 12,902 non-redundant SNPs, of which 9,493 were significantly associated with disorders of the immune system. Fisher’s exact test was used to calculate enrichment of immune trait-associated SNPs and non-immune trait-associated SNPs, against a background of common genetic variants (Common dbSnp153), in cell type-resolved peak sets or a control background set of peaks. Trait-specific enrichment analysis was performed using cell type-specific marker peaks (FDR < 0.05, log2FC > 0.25), with a background SNP set comprising all fine-mapped SNPs across all traits. Cell type-and tissue-specificity of accessibility at SNPs was determined by presence or absence of a scATAC peak in each cell type, with cell type clusters regrouped based on enrichment in tonsils or peripheral blood/bone marrow samples. Of the immune-related SNPs that overlapped with accessible chromatin peaks (1213, 12.8%), we subsequently identified 460 unique immune-linked SNPs that fell within 358 chromatin accessible regions for which a significant Peak2Gene link had been identified to at least one gene (P2G_Correlation > 0.4; FDR < 0.01). Mean normalized chromatin accessibility counts (scATAC) and RNA expression counts for linked genes (scRNA) for each cell type cluster were calculated and used for heatmap visualization while pyGenomeTracks was used to visualize grouped ATAC pseudobulk tracks (70). Linkage disequilibrium scores of top candidate SNPs were calculated using LDlink across all populations (71).

### Accession codes and data availability

Raw and processed data for this study are available at Gene Expression Omnibus under accession GSE165860.

## Author contributions

H.W.K., K.L.W., Z.S. and W.J.G. conceived the project and designed experiments. H.W.K., Z.S., A.S.K. and L.E.W. processed samples for single cell experiments. L.E.W performed CyTOF experiments and data analysis. H.W.K., K.L.W., and Z.S. performed scRNA and scATAC data analysis. C.L. provided guidance for analysis and interpretation of GWAS variants. R.C. and N.O. provided tonsillectomy surgical samples. H.W.K., K.L.W., Z.S. and W.J.G. wrote the manuscript with input from all authors. M.M.D, L.M.S., L.K.J. and W.J.G supervised the work.

## Supplementary Materials

Figure S1. Comparison of scRNA-seq and scATAC-seq broad immune cell clusters.

Figure S2. Comparison of RNA expression, cell surface protein expression and chromatin accessibility of key marker genes.

Figure S3. Age-related changes in tonsillar immune cell populations by scRNA-seq and CyTOF.

Figure S4. Tonsillar immune cell subset differential scATAC peaks.

Figure S5. Integrated bone marrow, blood and tonsil scRNA-seq and scATAC-seq markers.

Figure S6. Tonsil B cell-enriched gene expression markers compared to peripheral blood B cells.

Figure S7. Enrichment of fine-mapped autoimmune variants in immune cell subsets.

Figure S8. Genome snapshots of fine-mapped autoimmune variants at GZMB/GZMH, NKX2-3 and COTL1/KHLH36 loci.

Figure S9. Genome snapshots of fine-mapped autoimmune variants at KSR1/LGALS9 and TNFRSF1A/LTBR loci.

Figure S10. Genome snapshots of germinal center-associated cell type-specific regulatory activity at fine-mapped autoimmune variants at CD80, PRAG1 and SLC38A9/DDX4 loci.

Figure S11. Genome snapshots of germinal center-associated cell type-specific regulatory activity at fine-mapped autoimmune variants at VAV3 and DLEU2 loci.

Figure S12. Linkage disequilibrium scores for variants at IL21, IL21R and BCL6 loci.

Figure S13. Genome snapshots of fine-mapped autoimmune variants at ETS1 and IKZF3 loci. Figure S14. Genome snapshots of fine-mapped autoimmune variants at STAT4 and IRF8 loci.

Figure S15. Genomic landscape at HHEX and expression of KLF family transcription factors.

Table S1. CITE-seq antibody details.

Table S2. Broad resolution of tonsil immune cell subset scRNA-seq gene expression markers.

Table S3. High resolution B cell subset scRNA-seq gene expression markers.

Table S4. High resolution T cell subset scRNA-seq gene expression markers.

Table S5. Differential chromatin accessibility peaks from high resolution annotation of tonsil immune cell scATAC-seq.

Table S6. Differential chromatin accessibility gene scores from high resolution annotation of tonsil immune cell scATAC-seq.

Table S7. Gene expression markers for integrated tonsil, peripheral blood and bone marrow immune cell populations from scRNA-seq.

Table S8. Differential chromatin accessibility peak markers for integrated tonsil, peripheral blood and bone marrow immune cell populations from scATAC-seq.

Table S9. Peak2gene linkage annotation of fine-mapped SNPs found in chromatin accessibility peaks from integrated tonsil, peripheral blood and bone marrow scATAC-seq datasets.

---

**Figure S1.**
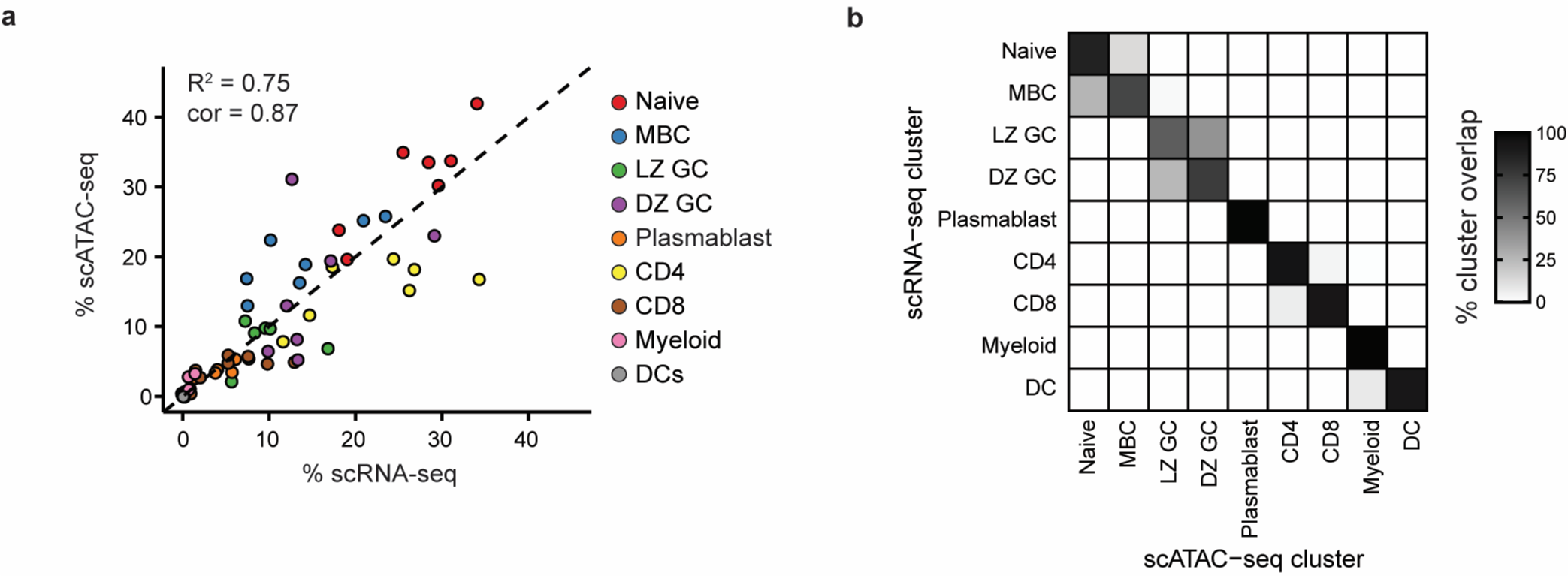
Comparison of scRNA-seq and scATAC-seq broad immune cell clusters. a) Correlation of relative cluster frequencies for scRNA-seq and scATAC-seq within each donor dataset. b) Confusion matrix depicting overlap between transferred scRNA-seq cluster identities to scATAC-seq clusters.

**Figure S2.**
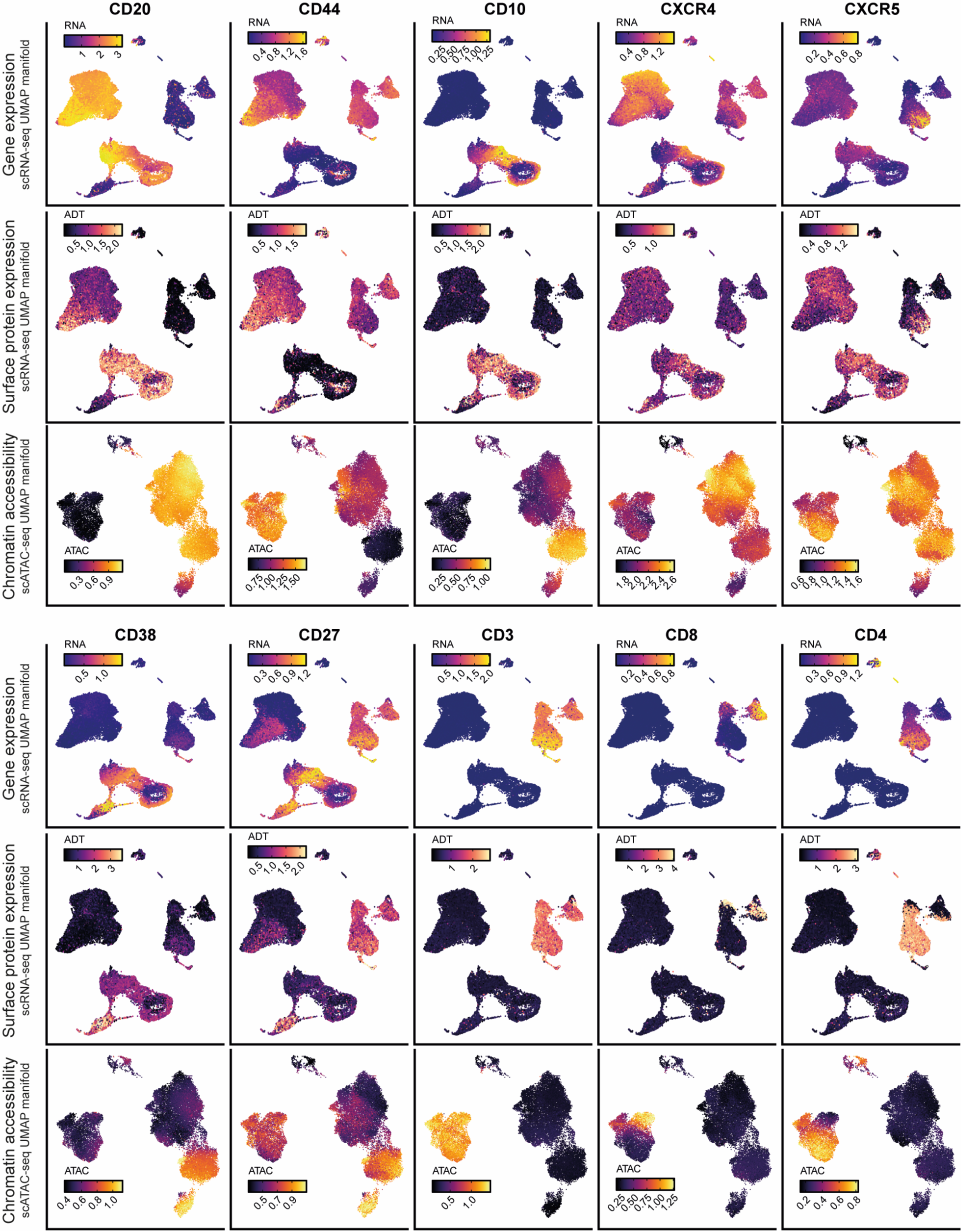
Comparison of RNA expression, cell surface protein expression and chromatin accessibility of marker genes. NA-seq gene expression (top rows), CITE-seq surface protein expression (middle rows) and chromatin accessibility gene es (bottom rows) for key marker genes. Gene expression and surface protein expression are visualized on the scRNA-seq P manifold and chromatin accessibility scores are visualized on the scATAC-seq UMAP manifold (see Figure 1b).

**Figure S3.**
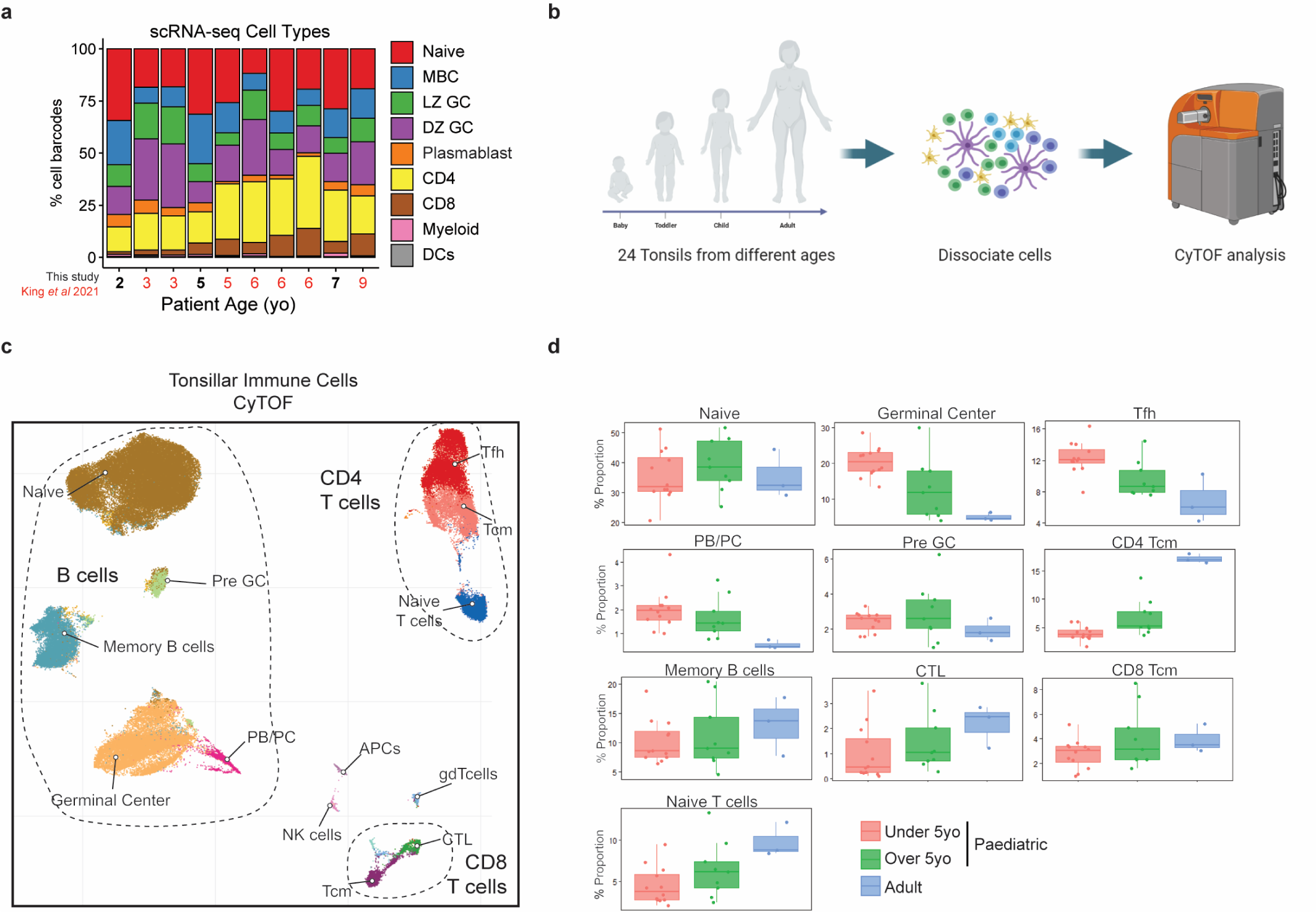
Age-related changes in tonsillar immune cell populations by scRNA-seq and CyTOF. a) Relative scRNA-seq cluster frequencies of different donors ordered by patient age. Additional tonsillar immune cell samples from King et al. (2021) are included (see red labels). b) Schematic of CyTOF analyses for age-related differences in tonsillar immune cell populations. c) UMAP visualization of CyTOF analysis of tonsillar immune cells with major immune cell populations (n=24). d) Quantitation of relative frequencies of immune cell subsets separated by patient age.

**Figure S4.**
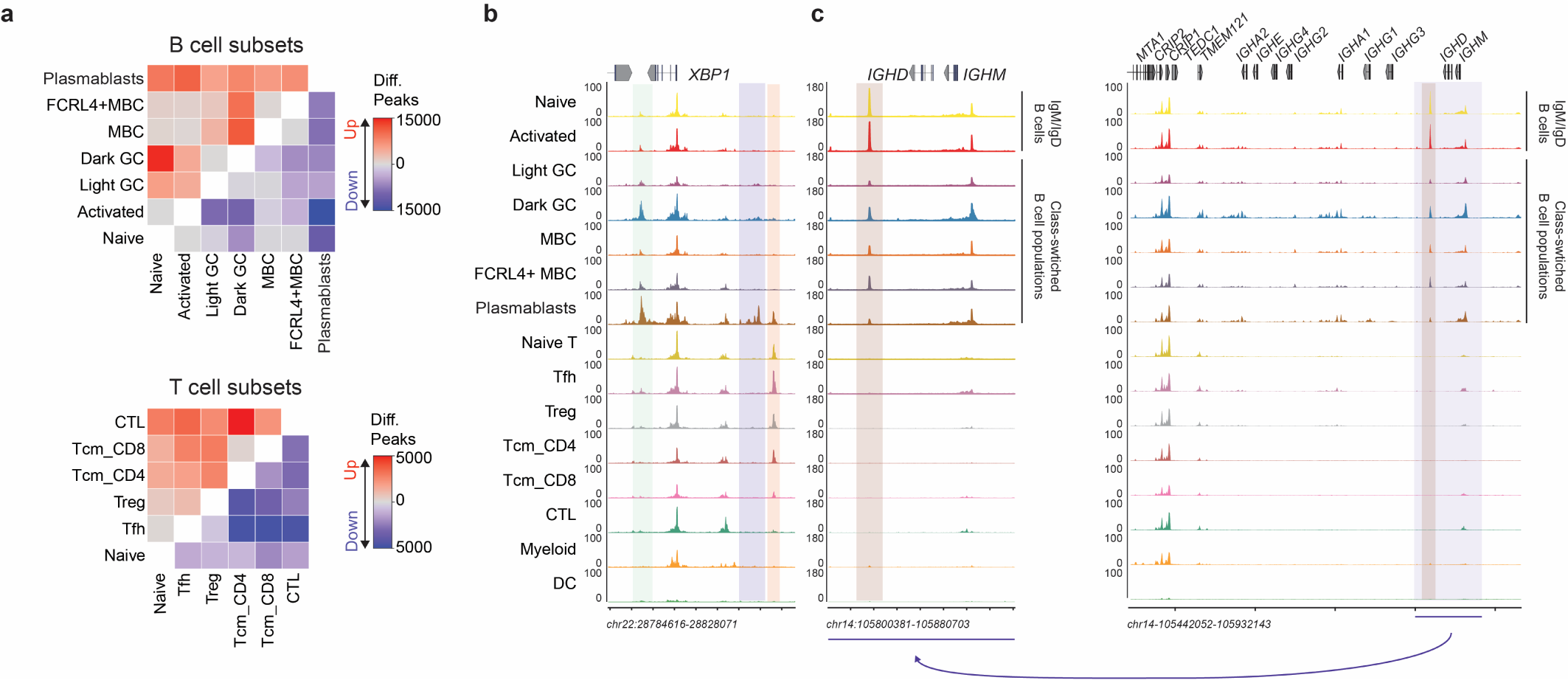
Tonsillar immune cell subset differential scATAC peaks. a) Differential peak analysis of B (top) and T (bottom) cell subsets comparing number of up-regulated and down-regulated chromatin accessibility regions. b) Example genome snapshot of XBP1 regulatory landscape. B cell-, plasmablast- and T cell-specific regulatory elements are highlighted. c) Genome snapshot of immunoglobulin heavy chain locus, including closer resolution of regulatory element downstream of IGHD and IGHM that is lost during class switch recombination (i.e. through deletional recombination).

**Figure S5.**
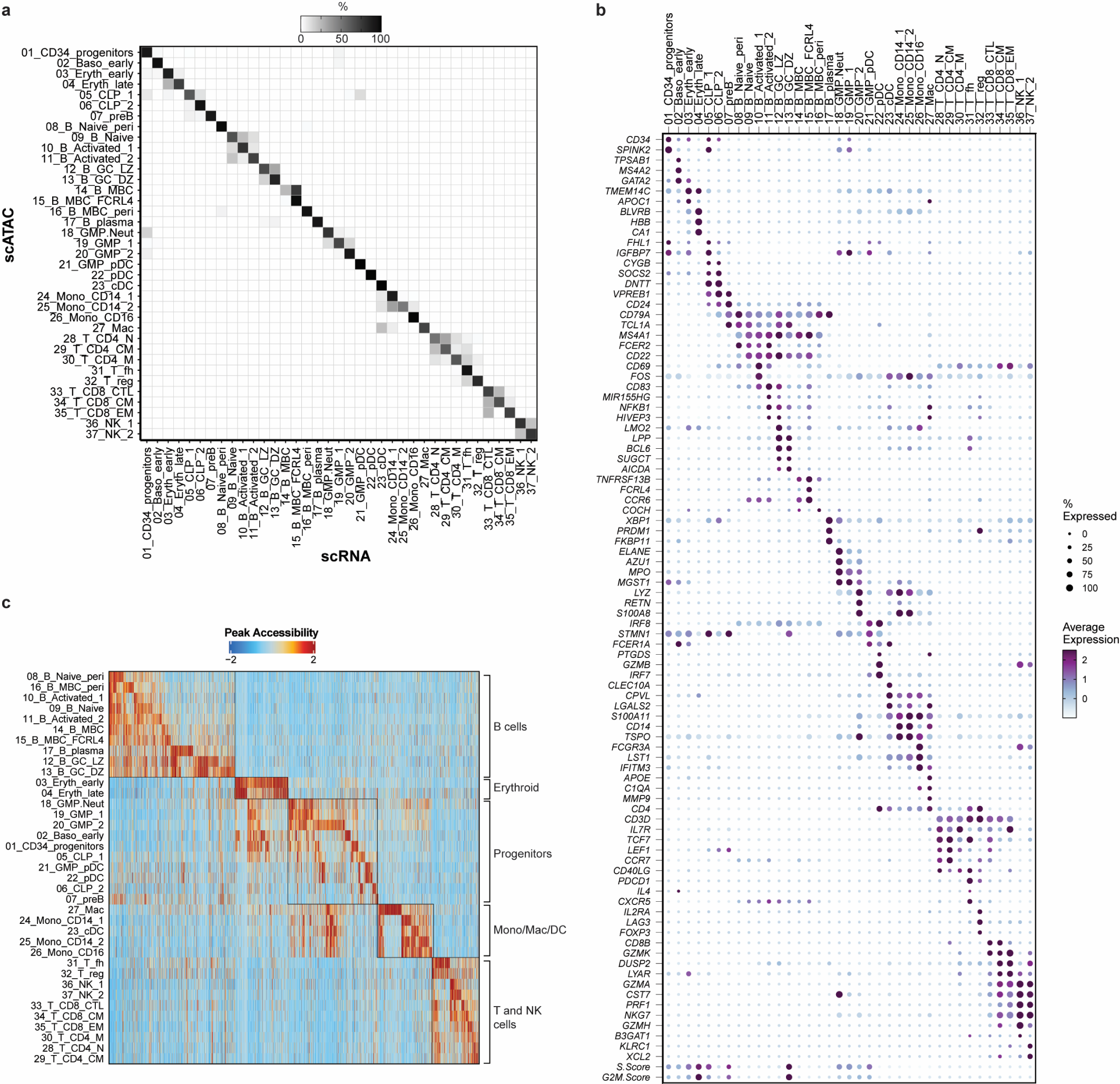
Integrated bone marrow, blood and tonsil scRNA-seq and scATAC-seq markers. a) Confusion matrix depicting overlap between transferred scRNA-seq cluster identities to scATAC-seq clusters. b) Expression of top marker genes for scRNA-seq clusters of integrated bone marrow, blood and tonsil dataset. c) Chromatin accessibility at cluster-specific peaks for scATAC-seq clusters of integrated bone marrow, blood and tonsil dataset.

**Figure S6.**
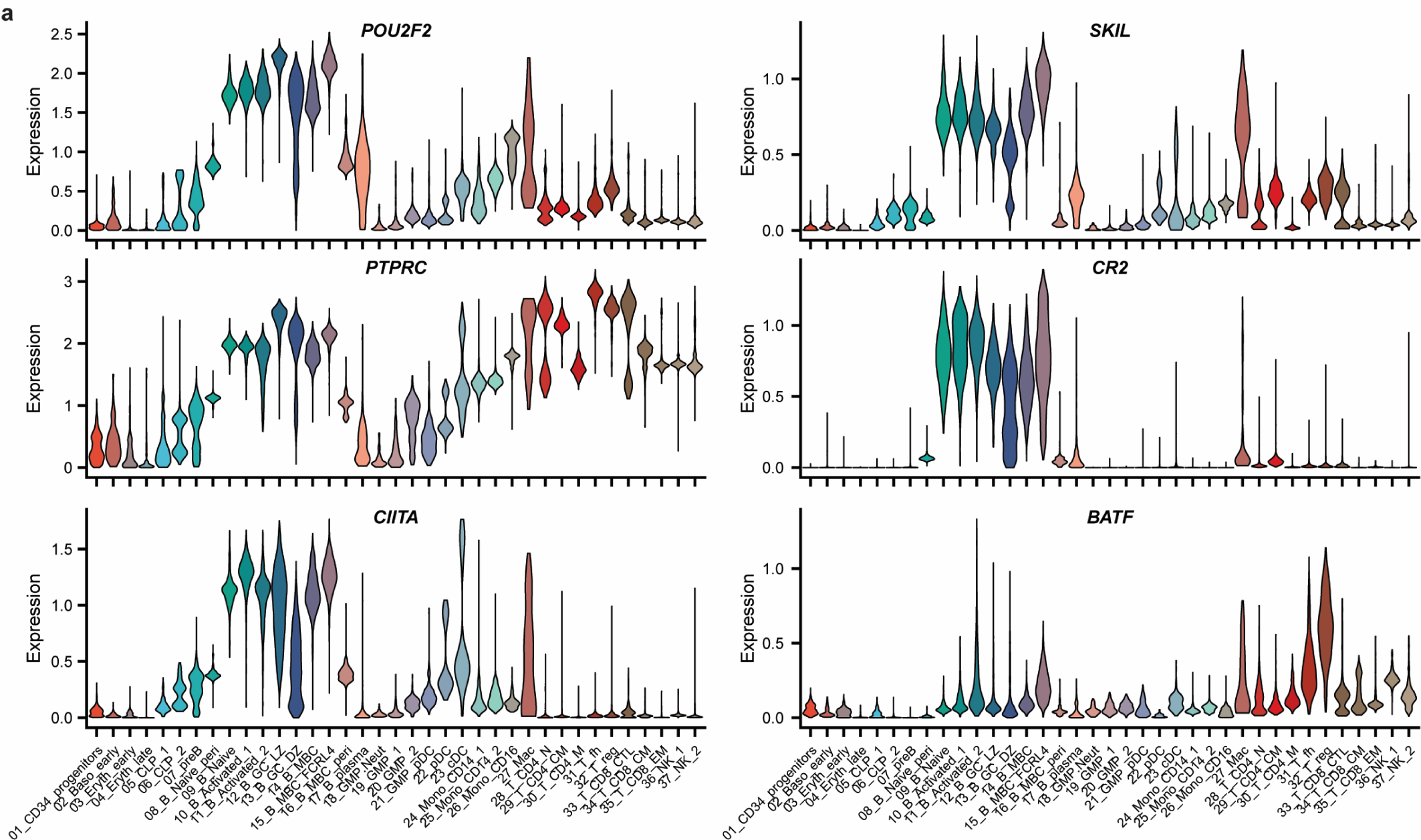
Tonsil B cell-enriched gene expression markers compared to peripheral blood B cells. a) Expression of genes significantly differentially expressed between tonsil-specific naive or memory B cell clusters compared to peripheral blood naive or memory B cell clusters.

**Figure S7.**
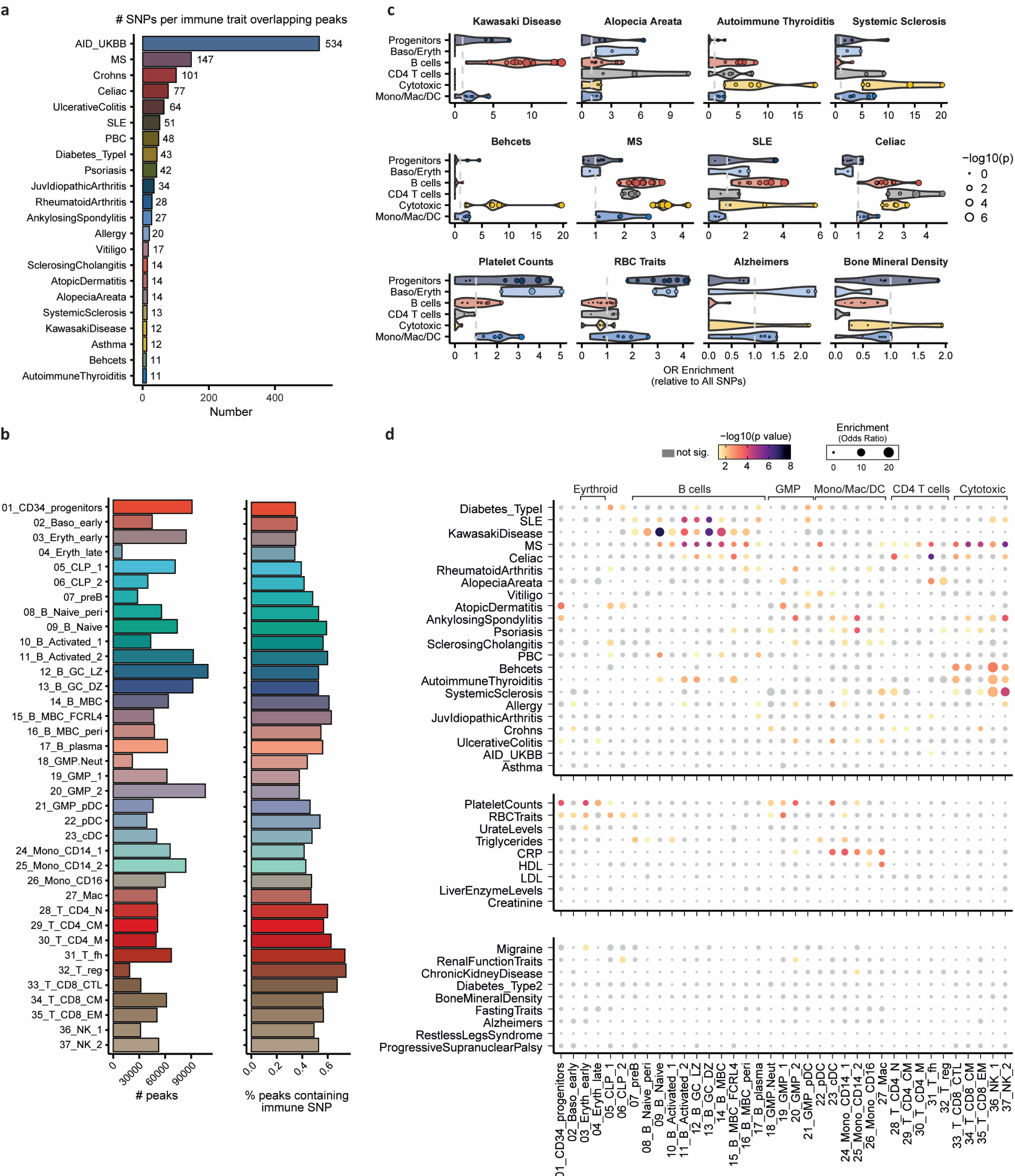
Enrichment of fine-mapped autoimmune variants in immune cell subsets. a) Number of fine-mapped SNPs per autoimmune trait that overlap with a chromatin accessibility peak in the integrated human bone marrow, peripheral blood and tonsil scATAC-seq atlas. AID_UKBB represents variants identified from fine­ mapping a combination of datasets from diverse autoimmune traits. AID; autoimmune disease. MS; multiple sclerosis. SLE; systemic lupus erythematosus. PBC; primary biliary cirrhosis b) Number of peaks identified in each scATAC-seq cell type cluster (left) and the percentage of those peaks that overlap with an autoimmune-associated SNP. c) Fisher enrichment test results for variants specific to selected traits in cell type-specific chromatin. Individual points represent single cell type clusters, separated into five broad lineages. Dot size reflects the level of significance of **enrichment.** d) Fisher enrichment test results for trait-specific variants in cell type-specific chromatin across all traits in fine-mapped resources analyzed. Dot size conveys enrichment (odds ratio) and color denotes significance of enrichment.

**Figure S8.**
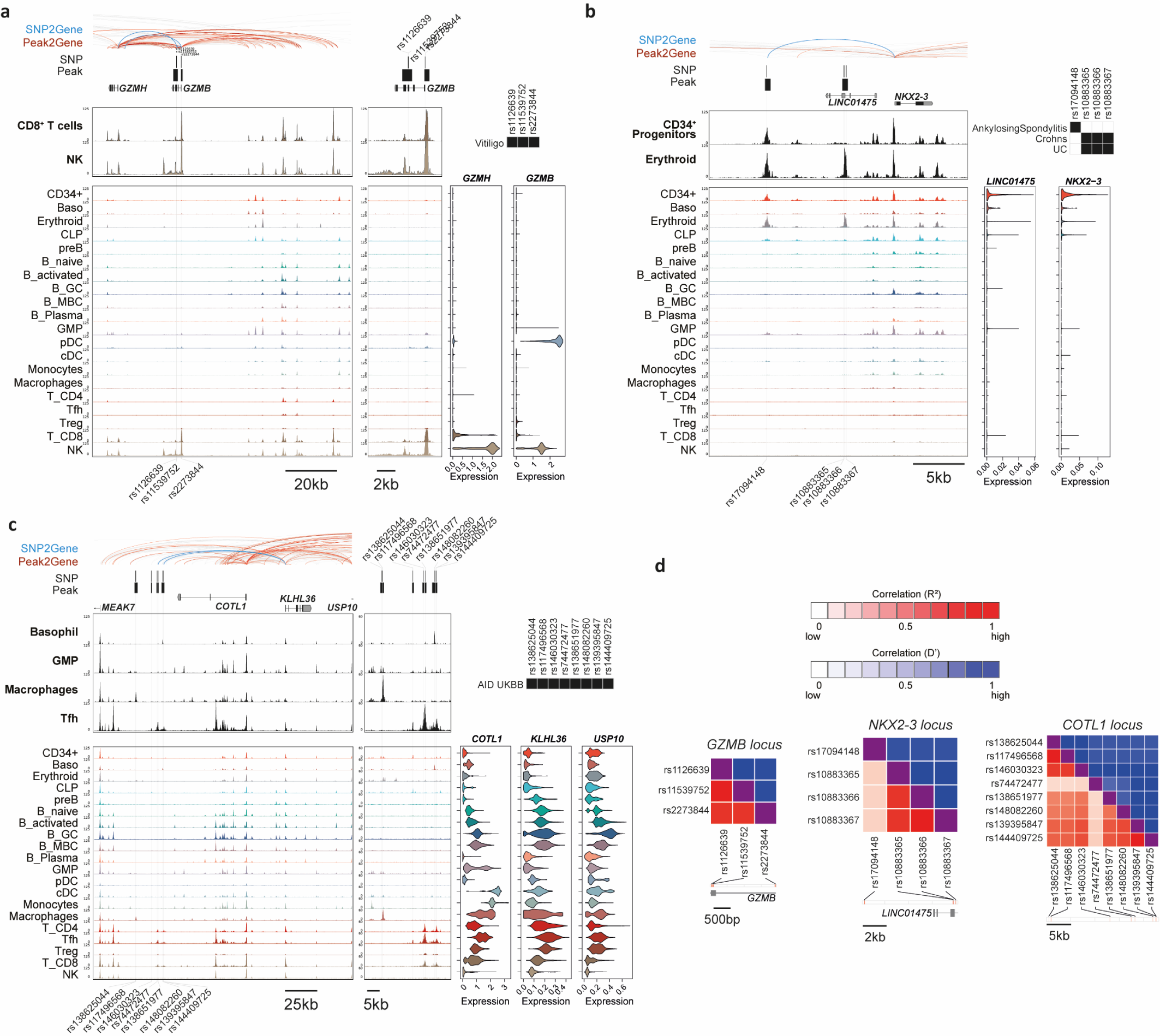
Genome snapshots of fine-mapped autoimmune variants at GZMB/GZMH, NKX2-3 and COTL1/KHLH36 loci. a) Genomic snapshot of fine-mapped autoimmune-associated GWAS variants at the GZMB/GZMH locus. Significant peak2gene linkages colored in red and significant links between SNPs and gene promoters (SNP2gene) in blue and bold. Significant associations between individual SNPs and autoimmune diseases are shown in black boxes and gene expression is shown as violin plots for matched populations in scATAC tracks. b) Same as A), at the NKX2-3 locus. UC; ulcerative colitis. c) Same as A), at the COTL1/KLHL36 locus. AID; autoimmune disease. d) Linkage disequilibrium heatmaps for SNPs at loci depicted in A-C. D′ denotes normalized linkage disequilibrium; R^2^ denotes Pearson coefficient of correlation.

**Figure S9.**
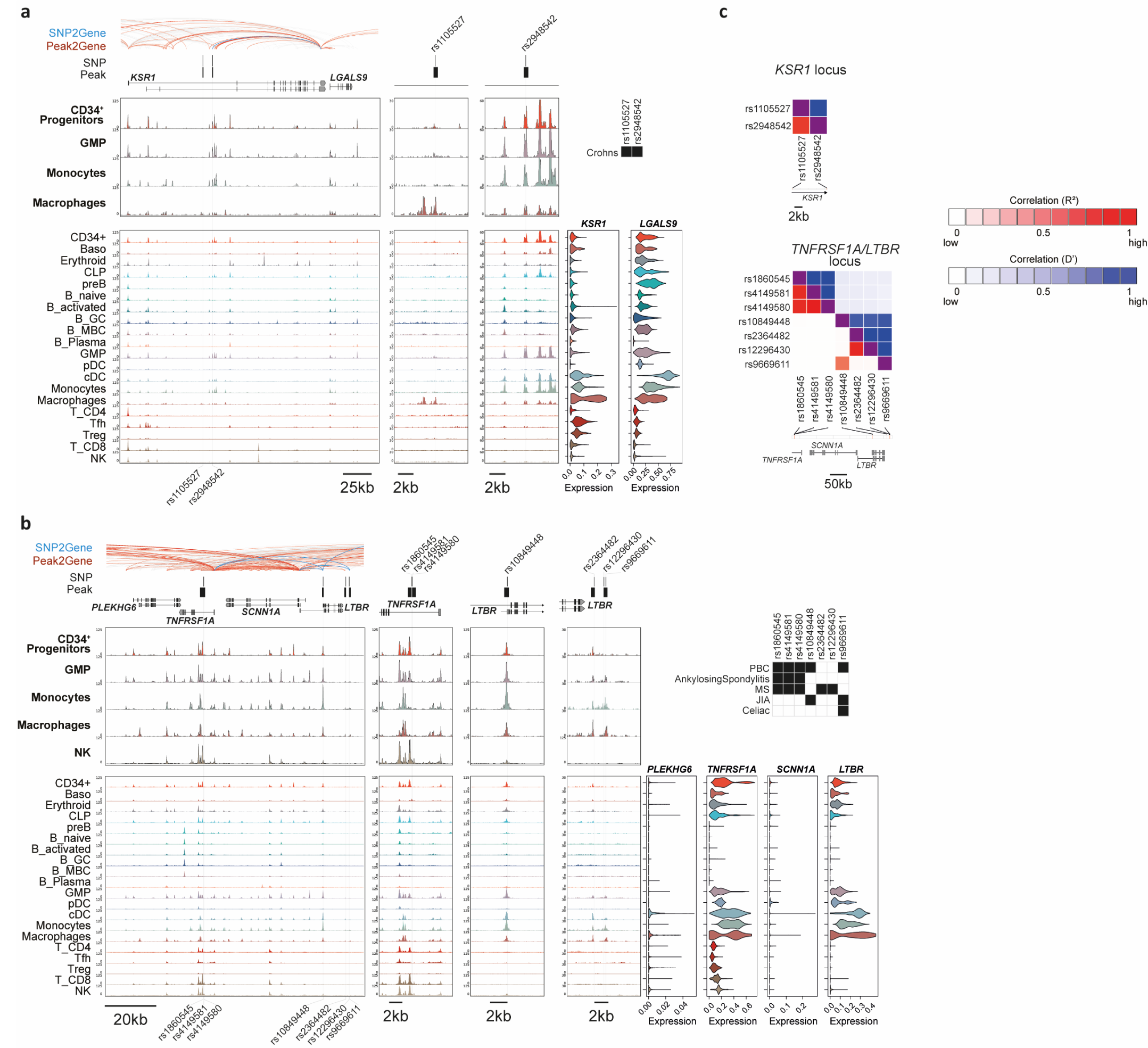
Genome snapshots of fine-mapped autoimmune variants at KSR1/LGALS9 and TNFRSF1A/LTBR loci. a) Genomic snapshot of fine-mapped autoimmune-associated GWAS variants at the KSR1/LGALS9 locus. Significant peak2gene linkages colored in red and significant links between SNPs and gene promoters (SNP2gene) in blue and bold. Significant associations between individual SNPs and autoimmune diseases are shown in black boxes and gene expression is shown as violin plots for matched populations in scATAC tracks. b) Same as A), at the TNFRSF1A/LTBR locus. PBC; primary biliary cirrhosis. MS; multiple sclerosis. JIA; juvenile idiopathic arthritis. c) Linkage disequilibrium heatmaps for SNPs at loci depicted in A and B. D′ denotes normalized linkage disequilibrium; R^2^ denotes Pearson coefficient of correlation.

**Figure S10.**
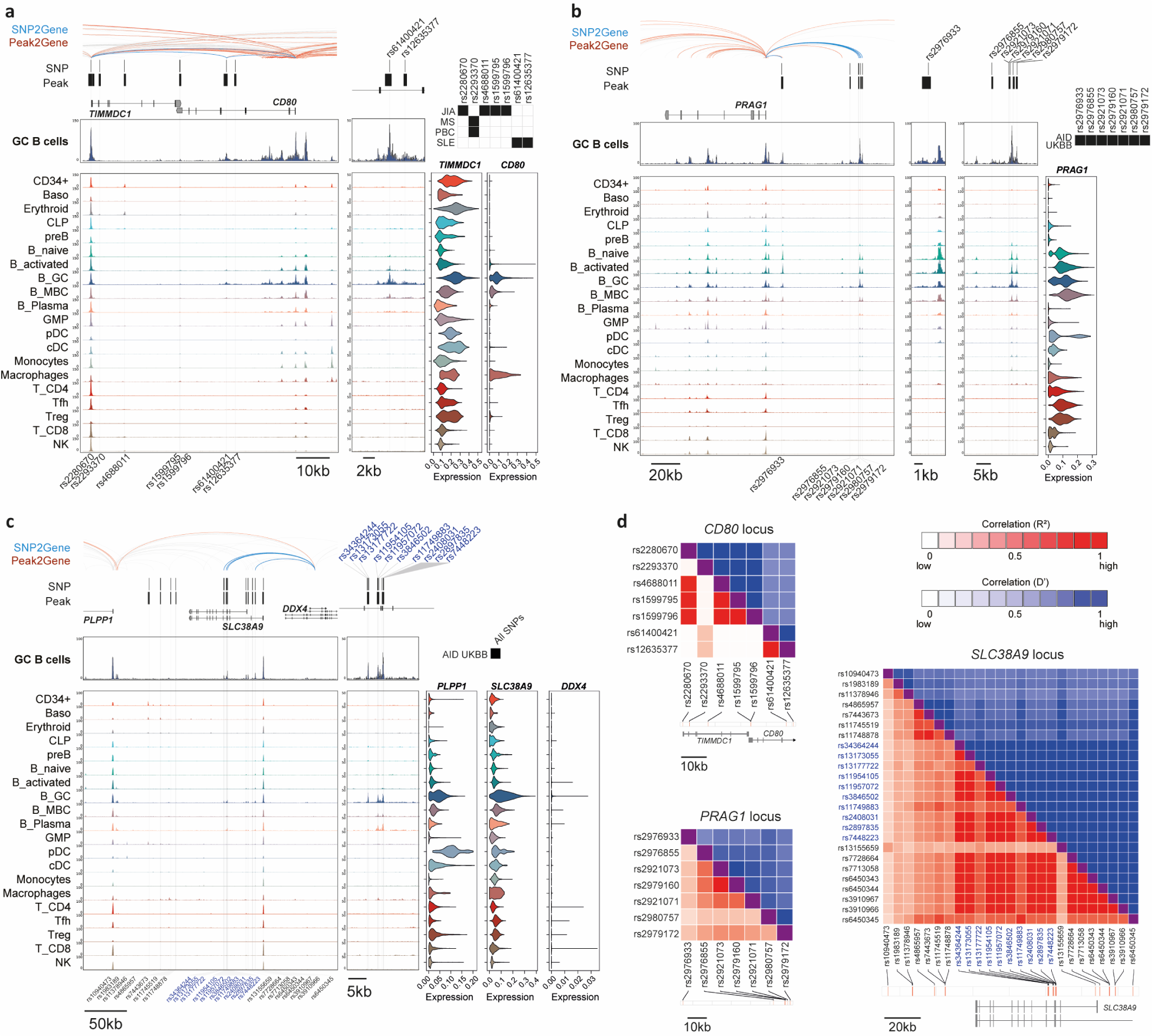
Genome snapshots of germinal center-associated cell type-specific regulatory activity at fine-mapped autoimmune variants at CD80, PRAG1 and SLC38A9/DDX4 loci. a) Genomic snapshot of fine-mapped autoimmune-associated GWAS variants at the CD80 locus. Significant peak2gene linkages colored in red and significant links between SNPs and gene promoters (SNP2gene) in blue and bold. Significant associations between individual SNPs and autoimmune diseases are shown in black boxes and gene expression is shown as violin plots for matched populations in scATAC tracks. PBC; primary biliary cirrhosis. MS; multiple sclerosis. JIA; juvenile idiopathic arthritis. SLE; systemic lupus erythematosus. b) Same as A), at the PRAG1 locus. AID; autoimmune disease. c) Same as A), at the SLC38A9/DDX4 locus. d) Linkage disequilibrium heatmaps for SNPs at loci depicted in A and B. D′ denotes normalized linkage disequilibrium; R^2^ denotes Pearson coefficient of correlation.

**Figure S11.**
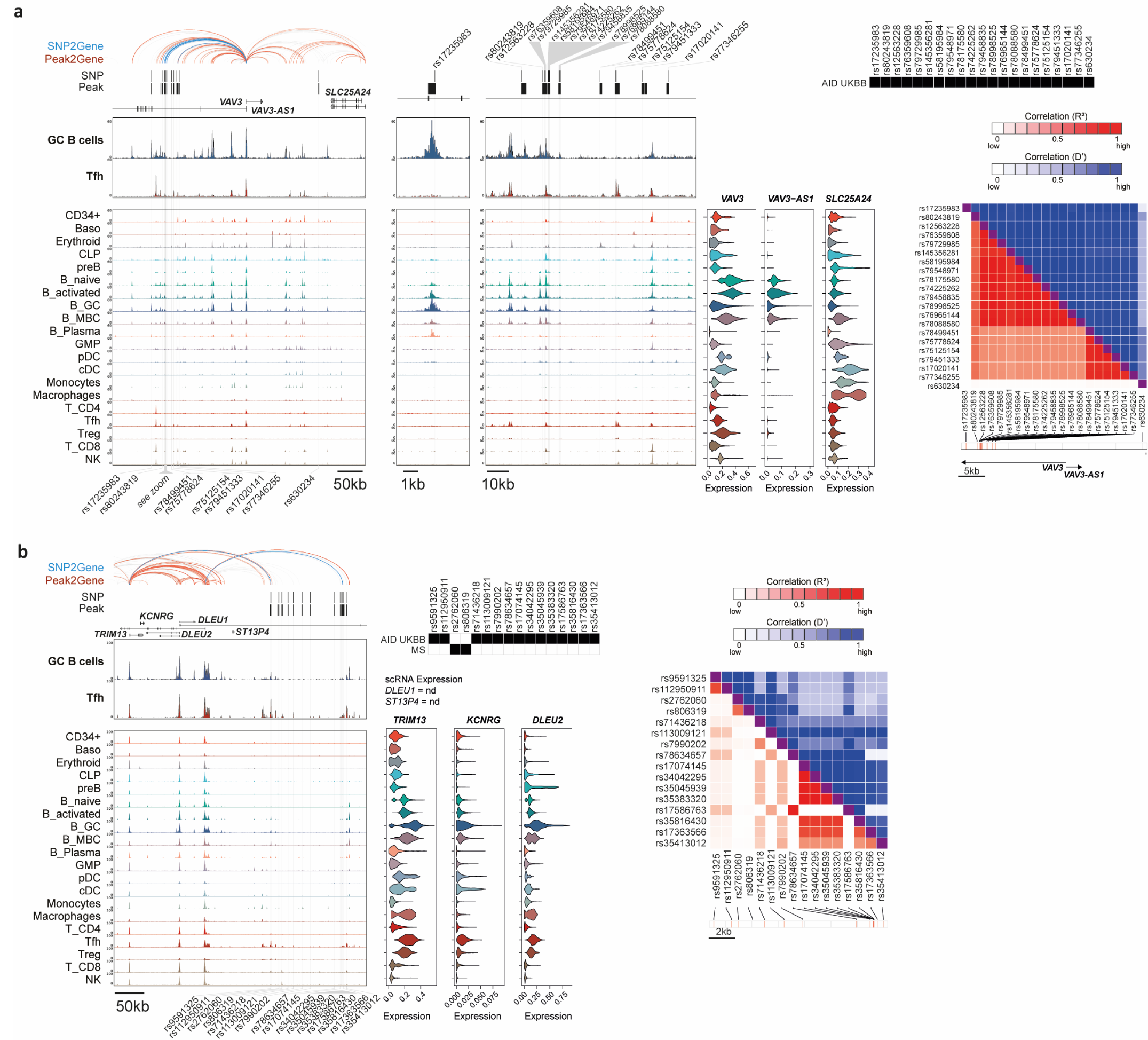
Genome snapshots of germinal center-associated cell type-specific regulatory activity at fine-mapped autoimmune variants at VAV3 and DLEU2 loci. a) Genomic snapshot of fine-mapped autoimmune-associated GWAS variants at the VAV3 locus. Significant peak2gene linkages colored in red and significant links between SNPs and gene promoters (SNP2gene) in blue and bold. Significant associations between individual SNPs and autoimmune diseases are shown in black boxes and gene expression is shown as violin plots for matched populations in scATAC tracks. Linkage disequilibrium heatmaps are also shown separately. D′ denotes normalized linkage disequilibrium; R^2^ denotes Pearson coefficient of correlation. AID; autoimmune disease. b) Same as A), at the DLEU2 locus. MS; multiple sclerosis.

**Figure S12.**
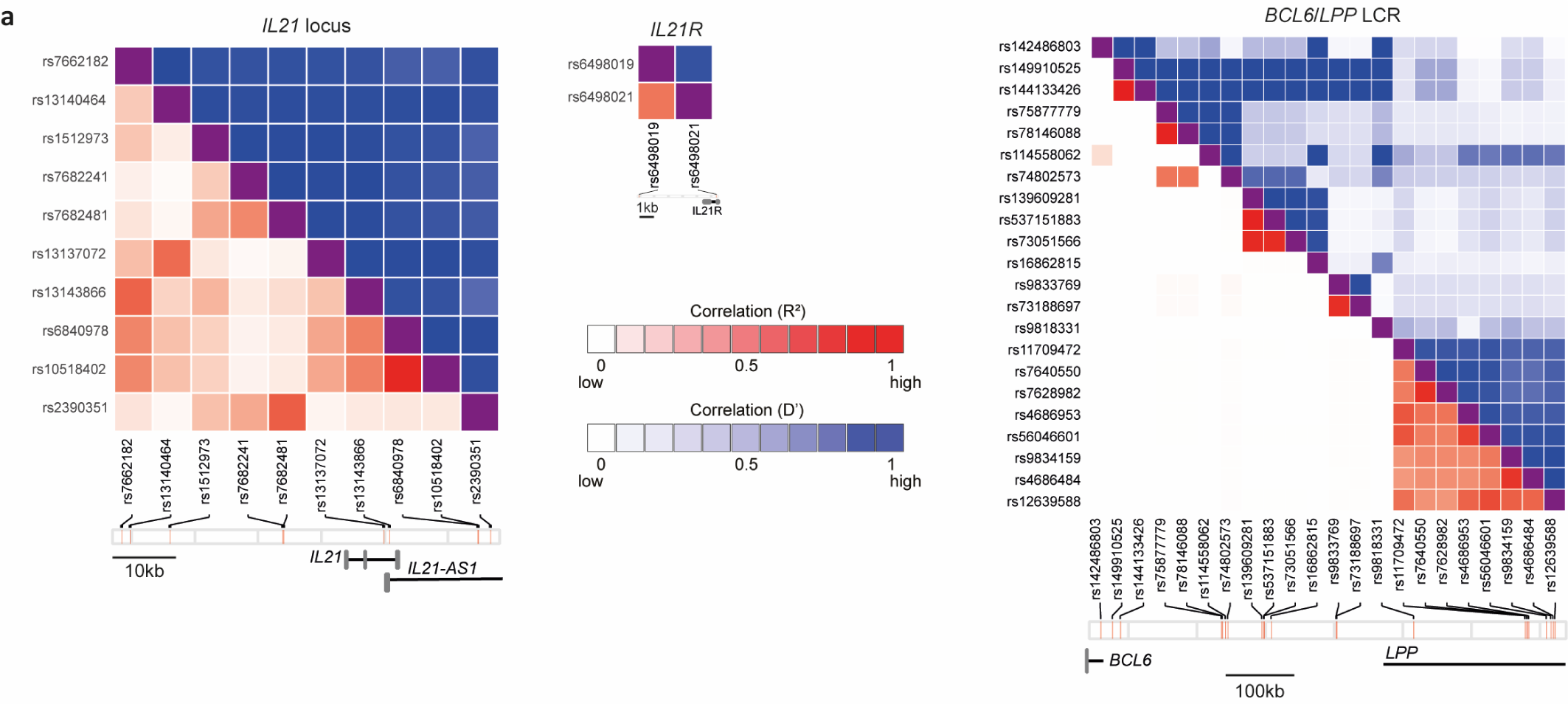
Linkage disequilibrium scores for variants at IL21, IL21R and BCL6 loci. a) Linkage disequilibrium heatmaps for SNPs at IL21, IL21R and BCL6/LPP loci depicted in Fig5. D′ denotes normalized linkage disequilibrium; R^2^ denotes Pearson coefficient of correlation.

**Figure S13.**
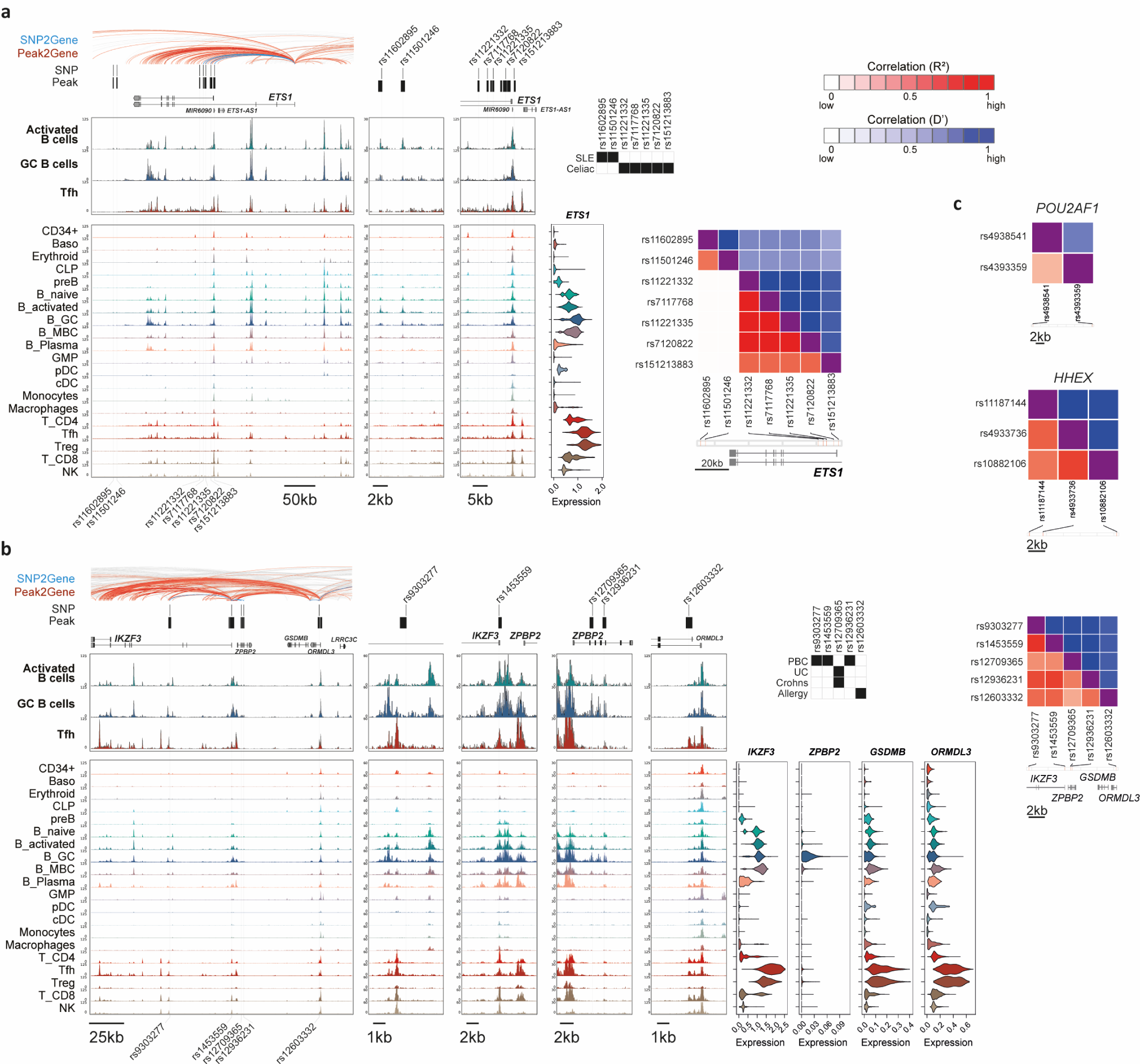
Genome snapshots of fine-mapped autoimmune variants at ETS1 and IKZF3 loci. a) Genomic snapshot of fine-mapped autoimmune-associated GWAS variants at the ETS1 locus. Significant peak2gene linkages colored in red and significant links between SNPs and gene promoters (SNP2gene) in blue and bold. Significant associations between individual SNPs and autoimmune diseases are shown in black boxes and gene expression is shown as violin plots for matched populations in scATAC tracks. Linkage disequilibrium heatmap is also shown. D′ denotes normalized linkage disequilibrium; R^2^ denotes Pearson coefficient of correlation. SLE; systemic lupus erythematosus. b) Same as A), at the IKZF3 locus. PBC; primary biliary cirrhosis. UC; ulcerative colitis. c) Linkage disequilibrium heatmaps for SNPs at POU2AF1 and HHEX loci depicted in Fig6. D′ denotes normalized linkage disequilibrium; R^2^ denotes Pearson coefficient of correlation.

**Figure S14.**
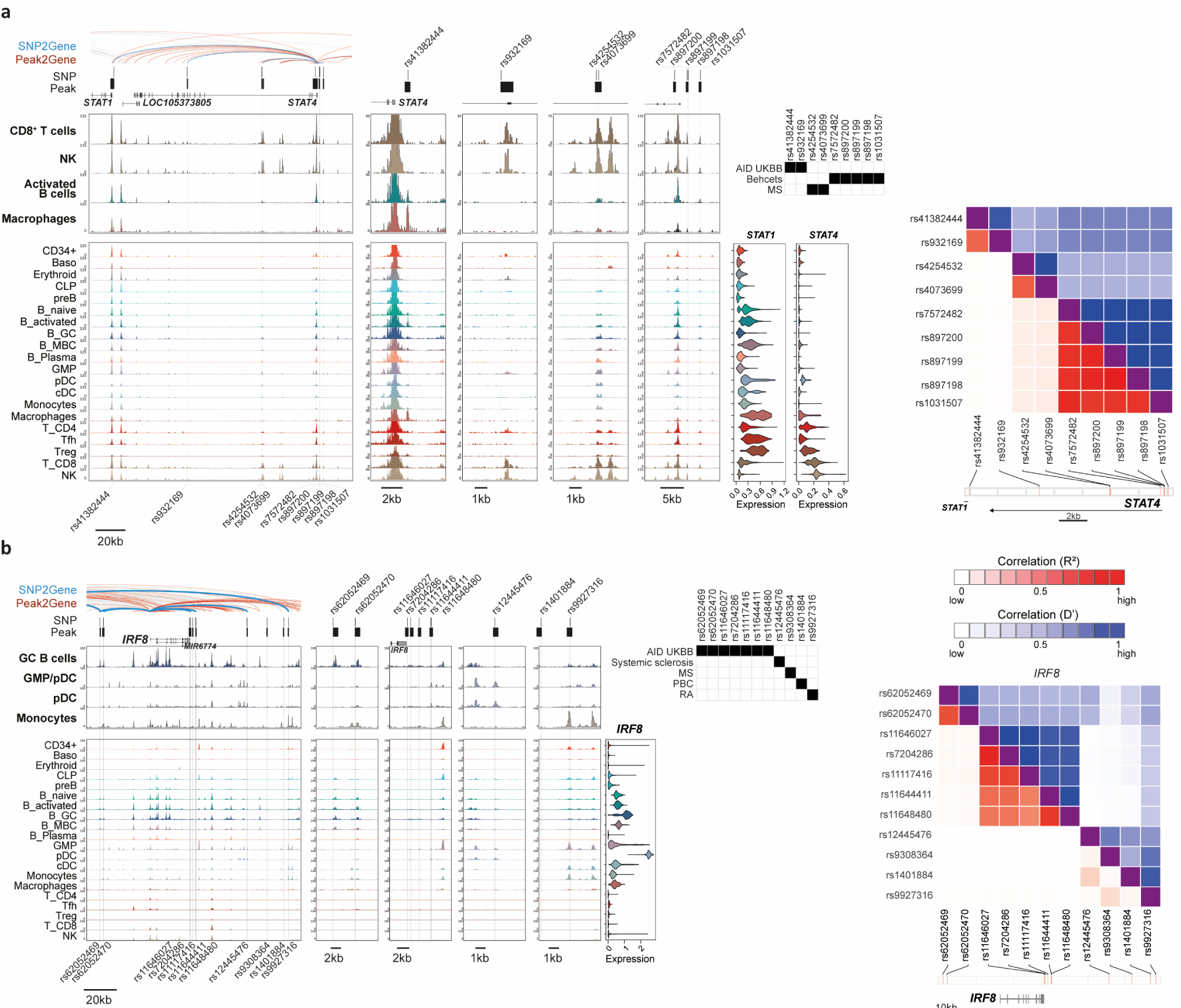
Genome snapshots of fine-mapped autoimmune variants at STAT4 and IRF8 loci. a) Genomic snapshot of fine-mapped autoimmune-associated GWAS variants at the STAT4 locus. Significant peak2gene linkages colored in red and significant links between SNPs and gene promoters (SNP2gene) in blue and bold. Significant associations between individual SNPs and autoimmune diseases are shown in black boxes and gene expression is shown as violin plots for matched populations in scATAC tracks. Linkage disequilibrium heatmap is also shown. D′ denotes normalized linkage disequilibrium; R^2^ denotes Pearson coefficient of correlation. AID; autoimmune disease. MS; multiple sclerosis. b) Same as A), at the IRF8 locus. PBC; primary biliary cirrhosis. RA; rheumatoid arthritis.

**Figure S15.**
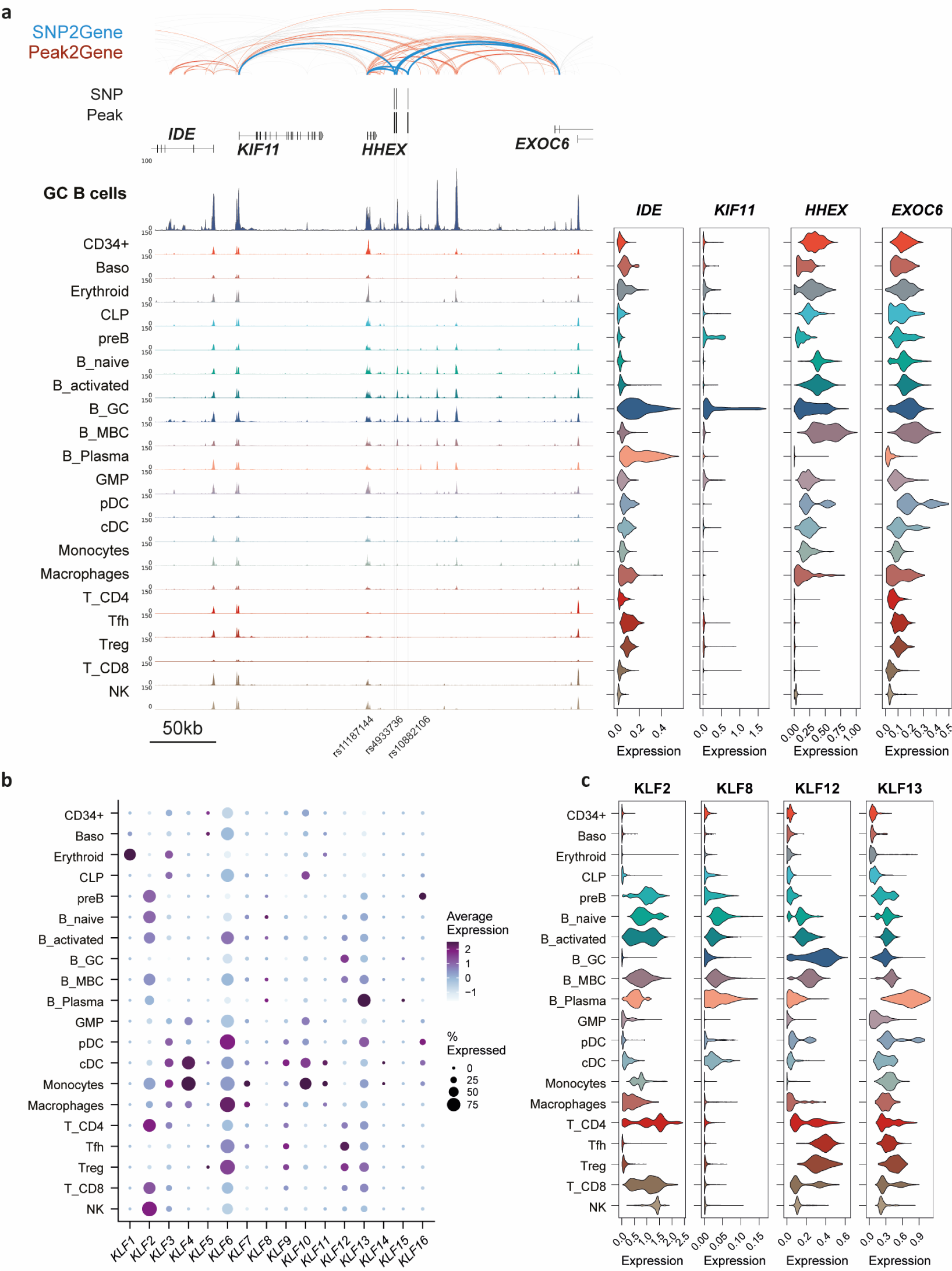
Genomic landscape at HHEX and expression of KLF family transcription factors. a) Broader view of the HHEX locus (see Fig6b), showing peak2gene links and gene expression of neighboring genes KIF11 and EXOC6. b) Mean expression of all KLF family transcription factors detected in scRNA-seq dataset. Dot size denotes percent of cluster in which gene is detected. c) Single-cell expression of KLF2, KLF8, KLF12 and KLF13, with highest expression in B cell subsets.

